# RNA-binding proteins identified by R-DeeP/TripepSVM are involved in heterocyst differentiation

**DOI:** 10.1101/2024.04.03.587981

**Authors:** Manuel Brenes-Álvarez, Halie R. Ropp, Dimitrios Papagiannidis, Clement Potel, Frank Stein, Mikhail M Savitski, Agustín Vioque, Alicia M. Muro-Pastor, Wolfgang R. Hess

**Affiliations:** Genetics and Experimental Bioinformatics, Faculty of Biology, University of Freiburg, 79104 Freiburg, Germany; European Molecular Biology Laboratory (EMBL), 69117 Heidelberg, Germany; Instituto de Bioquímica Vegetal y Fotosíntesis, Consejo Superior de Investigaciones Científicas and Universidad de Sevilla, 41092 Sevilla, Spain

## Abstract

RNA-binding proteins (RBPs) are central components of gene regulatory networks. The differentiation of heterocysts in filamentous cyanobacteria is an example of cell differentiation in prokaryotes. Although multiple non-coding transcripts are involved in this process, no RBPs have been implicated thus far. Here we used quantitative mass spectrometry to analyze the differential fractionation of RNA-protein complexes after RNase treatment in density gradients yielding 333 RNA-associated proteins, while a bioinformatic prediction yielded 311 RBP candidates in *Nostoc* sp. PCC 7120. We validated *in vivo* the RNA-binding capacity of 6 RBP candidates. Some participate in essential physiological aspects, such as photosynthesis (Alr2890), thylakoid biogenesis (Vipp1) or heterocyst differentiation (PrpA, PatU3), but their association with RNA was unknown. Validated RBPs Asl3888 and Alr1700 were not previously characterized. Alr1700 is an RBP with two OB-fold domains that is differentially expressed in heterocysts. Deletion of *alr1700* led to complete deregulation of the cell differentiation process, a striking increase in the number of heterocyst-like cells, and was ultimately lethal in the absence of combined nitrogen. These observations characterize this RBP as a master regulator of the heterocyst patterning and differentiation process, leading us to rename Alr1700 to PatR. The data can be accessed at https://sunshine.biologie.uni-freiburg.de/R-DeeP-Nostoc/.

## Introduction

Filamentous cyanobacteria are considered one of the oldest multicellular organisms on Earth^1^. Some of these organisms, such as *Nostoc* sp. PCC 7120 (hereafter *Nostoc),* have the ability to differentiate a specialized cell type devoted to nitrogen fixation, the heterocyst. Under nitrogen-replete conditions there are no heterocysts, but when the filaments encounter nitrogen starvation, the genetic program for their differentiation becomes induced in some vegetative cells following a semi-regular pattern. The developmental program takes about 24 hours to complete, providing an excellent experimental model for the analysis of cell differentiation in bacteria. Heterocysts are terminally differentiated, non-dividing cells dedicated to nitrogen fixation, and their morphology and physiology have evolved to create a microoxic environment that protects and supports nitrogenase activity^2^. During diazotrophic growth, vegetative cells continue to fix photosynthetic CO_2_, while heterocysts fix N_2_. Therefore, both types of cells must exchange metabolites to maintain the growth of the filament as a whole, providing one of the simplest examples of a true multicellular organism with two cell types cooperating together^3^.

NtcA and HetR have been recognized as the main transcriptional regulators of this differentiation process^4^. However, in striking parallel to more complex eukaryotic organisms, genome-wide analyses have suggested the involvement of hundreds of non-coding RNAs (antisense RNAs, asRNAs; small RNAs, sRNAs) in this process^5, 6^. Several sRNAs are regulated by the availability of nitrogen themselves and were designated <u>N</u>itrogen <u>s</u>tress induced <u>R</u>NAs (NsiR). Some have been characterized in more detail^7, 6, 8, 9^ and are involved in the reshaping of heterocyst metabolism^9^ or in the core regulation of the developmental process^8^. NsiR1 is expressed at a very early stage in the cells undergoing differentiation^10^ and inhibits, as an asRNA, the translation of the overlapping *hetF* mRNA^11^ encoding a protease crucially involved in heterocyst differentiation^12^. As a trans-acting sRNA, NsiR1 regulates the expression of *alr3234*^8^, encoding a HetP-like protein involved in the regulation of commitment to heterocyst differentiation^13^.

Regulation through prokaryotic non-coding RNAs frequently depends on RNA-binding proteins (RBPs) such as RNA chaperons or matchmakers, and extensive networks of post-transcriptional regulation have been elucidated^14^, but little is known about the interaction of RBPs and their functions in cyanobacteria. In fact, homologs to prominent RNA chaperones in proteobacteria, such as CsrA, ProQ, FinO or Hfq do not exist in cyanobacteria or do not bind RNA^15^.

RBPs play a central role in the physiology of cells. The ribosomes, the RnpB-RnpA (RNase P) complex, the signal recognition particle, the tmRNA-SmpB complex, or the vast diversity of CRISPR systems are well-known ribonucleoprotein complexes in prokaryotes. RBPs are key regulators of gene expression because they determine the fate of all RNA molecules in the cell^16^, including the regulation of mRNA stability or the spatial localization of mRNAs in the cell^17^. However, only a few pieces of information on the role of RBPs in the physiology of cyanobacteria are known. A family of RBPs containing an RNA recognition motif (RRM; Pfam: PF00076), similar to certain plant RBPs, was discovered in cyanobacteria almost 30 years ago^18^. Recently, they have been shown to be essential for the targeting of mRNAs encoding photosystem subunits near the thylakoid membrane, the site where photosynthetic proteins should be inserted^19^.

In this work, we have used R-DeeP^20^ to identify new RBPs expressed in *Nostoc* cell filaments that undergo heterocyst differentiation. We have identified 333 RNA-associated proteins that may be part of RNA-protein complexes. The *in vivo* validation of some of the candidates opens a new world of riboregulation in these organisms, as some RBPs have described functions not related to RNA metabolism, or were completely uncharacterized, such as Alr1700 (PatR), a protein with two OB-fold domains that affects the cell differentiation process in an intriguing way.

## Results

### *Nostoc* R-DeeP to fractionate multimeric complexes of a membrane-rich cyanobacterium

We cultivated triplicate cultures of *Nostoc* growing for 26 h in medium without a combined nitrogen source to obtain filaments containing vegetative cells, pro-heterocysts, and mature heterocysts for R-DeeP/GradR analysis^20, 21^. In this technique, all macromolecular complexes in the cell are fractionated in a density gradient followed by high-resolution proteomics of 19 fractions taken from each gradient. Gradients of RNase-treated samples were compared against those of untreated samples in triplicates. If a protein was part of an RNA-protein complex, the complex should be disassembled after RNase treatment and a shift of the protein position in the gradient should be observed (RNA-associated proteins) (Fig. 1a).

**Fig. 1.**
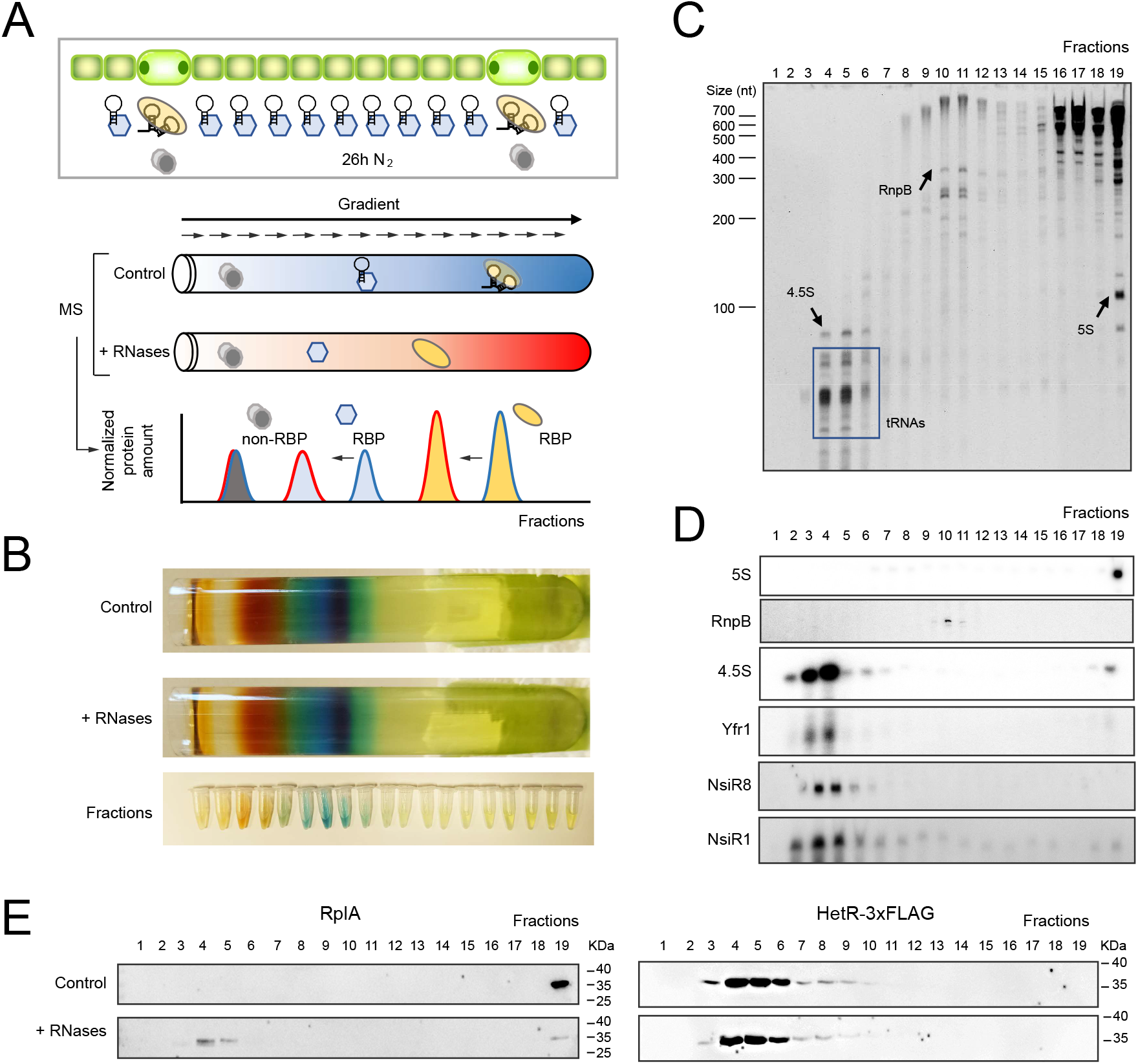
*Nostoc* R-DeeP experiment. **a** Overview: Lysates from heterocyst-containing filaments are fractionated in sucrose density gradients followed by MS proteomics. Gradients from RNase-treated and untreated samples are compared. An RNA-protein complex should dissociate after RNase treatment and a shift in the position of the respective proteins in the gradient should be observed. **b** Gradient tubes after centrifugation and fractions after elution in collection tubes. A representative pair of gradients is shown. The different colors correspond to abundant pigment-protein complexes, including orange carotenoids, blue phycobilins or green chlorophyll. **c** Separation of RNA from 19 fractions on a 10% urea-PAA gel. RNA was extracted from 300 µL of each fraction of one untreated gradient out of three. All extracted RNA was loaded onto the gel. The position of abundant RNA species is indicated for orientation. The position of size markers is indicated on the left. **d** Distribution of selected non-coding RNAs analyzed by Northern blot. RNA was extracted from 300 µL of each fraction of a representative untreated gradient, separated on a 10% urea-PAA gel and after blotting hybridized with radiolabeled probes for the indicated non-coding RNAs. **e** Distribution of RplA and HetR-3xFLAG along untreated and treated representative gradients. Western blot analyses were performed using 35 µL of each fraction separated on a 15% SDS-PAA gel and antisera against RplA or the FLAG epitope. The position of size markers (in kilodaltons) is indicated on the right. The results from one of three gradient pairs are displayed.

A critical step in the success of the approach is the preservation of the native protein-protein and RNA-protein complexes. Since cyanobacteria, as photosynthetic organisms, are rich in pigments, the fractionation of their macromolecular complexes resulted in colorful gradients reflecting the abundance of large photosynthetic complexes (Fig. 1b). We used 10-40% sucrose gradients, which have a resolution between 0 and 900 KDa. If the *Nostoc* proteome was completely disassembled into monomeric proteins, no proteins should be detected in fractions beyond fraction 5 (Supplementary Fig. 1a). However, proteins were measured in higher fractions (Supplementary Fig. 1b) and protein SDS-PAGE gels stained with Coomassie blue showed highly expressed protein complexes, such as RuBisCO or glutamine synthetase, appearing in higher fractions. Furthermore, the overall distribution of proteins did not appear to be affected by RNase treatment, excluding unspecific effects of treatment (Supplementary Fig. 1c-d). In order to test the integrity of the RNA in the untreated samples, RNA was extracted from the different fractions and separated in denaturing urea-polyacrylamide (PAA) gels. Several intact highly expressed RNAs were directly identified (Fig. 1c). We analyzed by Northern blot hybridization the distribution of 6 previously known non-coding RNAs; 5S rRNA, RnpB, 4.5S, Yfr1^7^, NsiR8^6^ and NsiR1^8^. The position of an RNA in the gradients did not depend on its size, but rather on its interaction with large protein complexes. This fact can be observed in the different distributions of the approximately equal-sized RNAs 4.5S (signal recognition particle RNA) and 5S rRNA (Fig. 1d). The detection of NsiR1 and NsiR8, two sRNAs expressed only in heterocysts, confirmed the capacity of detecting putative heterocyst-specific ribonucleoprotein complexes. This was a relevant aspect regarding the sensitivity of the analysis, because after heterocyst differentiation, there is on average only one heterocyst per 15 vegetative cells. Finally, as a positive control, the change in the position of the ribosomal RBP RplA after RNase treatment was tested by Western Blot (Fig. 1e). As a negative control, we generated a strain expressing the master transcription factor of heterocyst differentiation HetR^22^ fused to a 3xFLAG epitope tag under control of its native promoter. We prepared gradients with this strain following the same procedure as for the WT. The absence of a shift for HetR-3xFLAG confirmed the specificity of our approach (Fig. 1e).

Tandem mass tags (TMT)^23^ were used for the parallel detection and relative quantification of proteins by mass spectrometry (MS). We pooled fractions 1 and 2, thus 18 samples were measured per gradient. These measurements identified 2,638 proteins in at least one pair of control-RNase-treated gradients, which represent 44% of the *Nostoc* proteome. Forty of the 139 proteins (29%) were previously defined as being expressed exclusively in heterocysts^5^. The overall in-gradient distribution of protein profiles showed correlation scores of 0.92 to 0.94 between the replicates, and thus confirmed good reproducibility (Supplementary Fig. 2). To compare the distribution of proteins with different expression levels, an in-gradient normalization was performed, so that the sum of each protein along a gradient was set to 100%. The overall quality of the approach can be observed in the co-sedimentation of proteins belonging to essential ribonucleoprotein complexes such as 50S and 30S ribosomal subunits, RNA polymerase, or one of the *Nostoc* CRISPR systems*^24^* (Fig. 2a-d). To solubilize membrane proteins, we used n-dodecyl β-D-maltoside, a mild detergent commonly used to study photosynthetic membrane complexes^25^. The quality of our approach was confirmed by co-sedimentation of components from large membrane complexes involved in the bioenergetics of cyanobacteria, such as the PSII core, the PSI core, ATP synthase, NADH dehydrogenase, succinate dehydrogenase, cytochrome b_6_f or terminal oxidases, (Fig. 2e-l). The different oligomeric states of PSII and PSI can be observed in our data (Fig. 2e). PSII is mainly isolated in the monomeric state, whereas the PSI distribution is diffuse due to its capacity to form dimers and tetramers^26^. Different co-sedimentation profiles were observed for phycobilisomes, the major light harvesting system in red algae and cyanobacteria^27^, although the dissociation of phycobilisomes into subcomplexes is a biological response to nitrogen deprivation^28^.

**Fig. 2.**
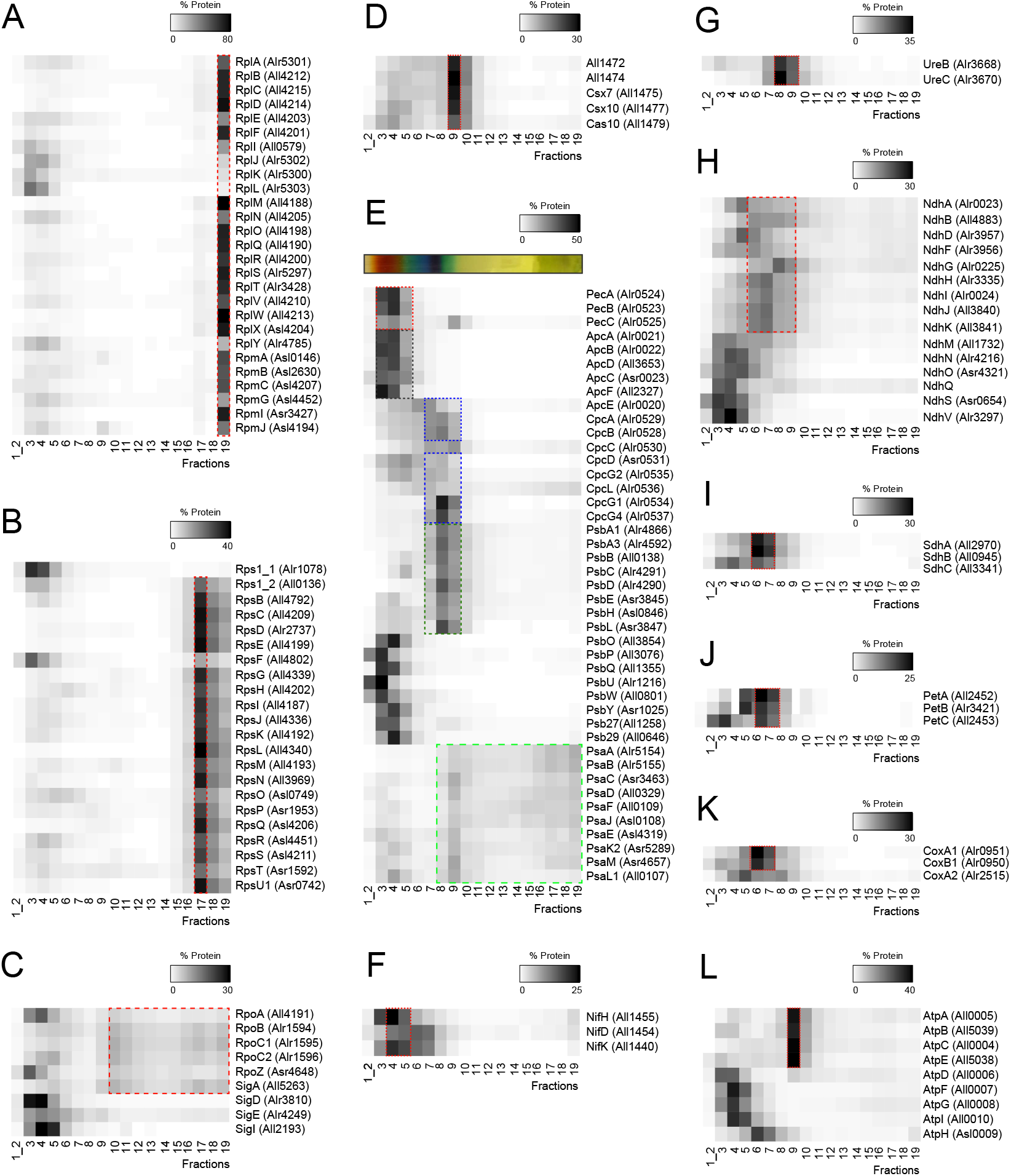
In-gradient distribution of proteins that are part of large macromolecular complexes in *Nostoc*. Heatmap representation of the mean normalized abundance of each detected protein in three untreated gradients. The sum of a protein abundance along the gradient is normalized to 100%. Proteins that are part of RNA-protein complexes are shown as follows: **a** 50S ribosomal subunit, **b** 30 ribosomal subunit, **c** RNA polymerase, **d** Subtype III-D CRISPR complex. Proteins that belong to multiprotein complexes involved in photosynthesis, nitrogen metabolism or the respiratory chain are also shown: **e** Photosynthetic complexes, **f** Nitrogenase, **g** Urease, **h** NADH dehydrogenase, **i** Succinate dehydrogenase, **j** Cytochrome b_6_f, **k** Terminal respiratory oxidases, **l** ATP synthase. Components of each complex with similar distributions are framed in red. In panel (**e**), a different color palette for the frames is used and a photo of the gradients is displayed above the heatmap to show the colors of the different fractions. Phycobilisome components containing phycoerythrocyanin, allophycocyanin or phycocyanin are framed in orange, black and blue, respectively. Co-sedimentation of the PSII and PSI core is also highlighted in dark green and light green, respectively.

The data allowed the identification of heterocyst-specific complexes, such as the nitrogenase core (Fig. 2f). Different combinations of core complexes with proteins present in all cells or specific to heterocysts were also observed. For example, two types of sigma factors co-sedimented with RNA-polymerase; the housekeeping sigma factor SigA, expressed in all cells, and the heterocyst-specific SigE factor^29^ (Fig. 2c). Another example is the different distribution of CoxB1 and CoxA1, components of a cytochrome *c* terminal oxidase that is expressed in vegetative cells, compared to the distribution of CoxA2, a component of a cytochrome *c* terminal oxidase only expressed in heterocysts^30^ (Fig. 2k). Normalized data for each protein can be found in Supplementary Data 1-2. We also provide a comprehensive resource, available at https://sunshine.biologie.uni-freiburg.de/R-DeeP-Nostoc/to scan for all proteins detected in this dataset.

According to NCBI, 27% of the *Nostoc* coding sequences are still annotated as encoding hypothetical proteins or proteins with domains of unknown function (16/03/2023). The co-sedimentation of unknown proteins with well-studied ribonucleoprotein complexes could aid in the detection of novel components. For this reason, we performed a soft clustering analysis on the protein profiles using the mean of the triplicates for the control gradients (Supplementary Fig. 3). We grouped the proteins into 19 groups. The correct distribution in the PCA plot (Supplementary Fig. 3a) and an average silhouette of 0.34 confirmed the good clustering of the data. Unknown proteins in clusters 15-19 coincided with a low number of identified complexes. For example, cluster 18 contains only proteins that are part of the 30S ribosomal subunit plus two proteins not previously associated with the ribosome, Alr0189 (a kinase) and Asl3888 (an unknown protein that is discussed below). The classification of proteins into different clusters can be found in Supplementary Data 3.

### Identification of new RNA-binding proteins

We used a differential expression-like approach for the analysis of the R-DeeP data. We compared the accumulation of each protein in each fraction in triplicate using the *limma* package^31^. Proteins with a fold change ≥ 2 and fdr ≤ 0.05 in at least one fraction were considered hits, while proteins iw th a fold change ≥ 1.5 and fdr ≤ 0.02 ere considered candidates. A total of 333 proteins had a significant change in at least one fraction and were considered RNA-associated proteins (Fig. 3a and see Supplementary Data 4 or https://sunshine.biologie.uni-freiburg.de/R-DeeP-Nostoc/ for the full list of proteins, fold changes, and fdr values of each fraction). The GO-term for “RNA-binding” (GO:0003723) had a signicant enrichment in the 333 RNA-associated proteins (q-value = 2.25e-10) and 34 of the RNA-associated proteins were previously annotated as RBPs. These proteins share a significantly higher isoelectric point than the shifting proteins not previously characterized as RBPs (Fig. 3b). A heatmap visualizing the observed shifts for the RNA-associated proteins showed an unexpected pattern (Fig. 3a). Half of the proteins migrated, as expected, to lower fractions after RNase treatment (left shift), but the other half of the proteins migrated to higher fractions after treatment (right shift). Hierarchical clustering analysis of the log2 fold changes of the 333 RNA-associated proteins detected two main clusters (Supplementary Data 5), which mainly resembled the right- or left-shifting behavior (Fig. 3a).

**Fig. 3.**
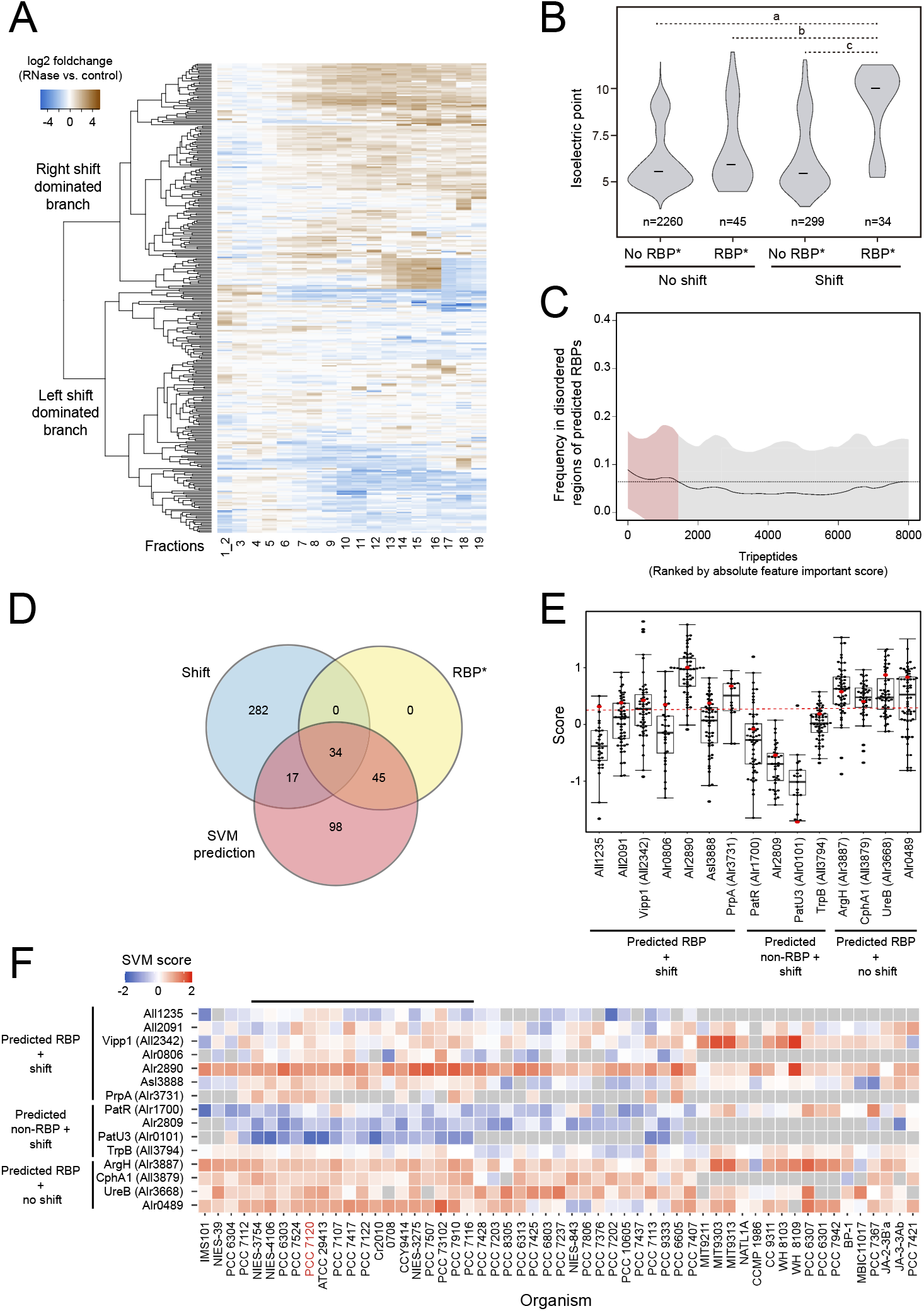
Bioinformatic analysis of *Nostoc* R-DeeP data. **a** Heatmap representation of log2 fold change data of shifting proteins. The log2 fold change (control vs. treated samples) of the 333 proteins with at least one difference (fold change ≥ 1.5 and fdr ≤ 0.02) in one of the fractions is shown. Blue shows the fractions in which a protein disappeared after RNase treatment, while brown shows the fractions to which it shifted after RNase treatment. **b** Violin plots showing the distribution of isoelectric points of the proteins identified by MS. The proteins that shift are those shown in (**a**). Proteins associated with “RNA binding” GO:0003723 in QuickGO are considered to be previously known *RBPs. The bar indicates the median in each group. The number of proteins belonging to each group is also shown. Statistically significant differences between groups (p-values < 0.01) by two-sided Wilcoxon test are shown with dashed lines (p-values; a=1.11x10^-13^, b=2.958x10^-6^ and c= 2.98x10^-11^). **c** Prevalence of tripeptides in structurally disordered regions of *Nostoc* predicted RBPs. The frequency of each tripeptide in structurally disordered regions (y-axis) is plotted, ranked by the absolute k-mer importance score of each tripeptide (x-axis). The resulting curve is smoothed using LOESS regression (span=0.2). The standard deviation is indicated by the shading. The dashed line marks the value of the 80% quantile of the smooth LOESS regression values. **d** Venn diagram showing the overlap between shifting proteins in our R-DeeP data, proteins annotated as ‘RNA-binding’ in QuickGO, and proteins predicted to be RBPs based on our modified TriPepSVM approach. **e** SVM score boxplot of selected proteins included in the groups shown in the Venn diagram of (**d**). Red dots show the SVM scores of *Nostoc* proteins. Black dots show the SVM scores of the cyanobacterial homologs. The red dashed line indicates the threshold to be considered an RBP. Proteins with a score greater than 0.25 are predicted to be RBPs. **f** SVM score heatmap of the proteins shown in (**e**). The unfound homologs of a protein are shown in grey. Higher SVM scores are shown in red, indicating a higher predicted ability to bind RNA. The horizontal black line encompasses the heterocyst-forming clade. Complete names of the displayed genomes can be found in Supplementary Table 2.

The shift of a protein in the gradients does not imply that the protein was binding RNA itself. Therefore, we implemented a modified version of TriPepSVM^32^, a support vector machine (SVM) approach that predicts RBPs based on tripeptide frequencies. As training data, we used proteins from 10 widely different cyanobacteria plus the well-annotated proteomes of *Escherichia coli* and *Salmonella typhimurium* (see Supplementary Table 1). A 10 k-fold cross validation nested on a grid search was used for the optimization of the best parameters (Supplementary Fig. 4a). In addition, a restrictive cutoff of 0.25 was selected to improve the specificity and clearly separated the RBPs from DNA- or nucleotide-binding proteins (Supplementary Fig. 4b). The prediction of RBPs using the entire *Nostoc* proteome yielded 311 candidates (Supplementary Data 6). We observed a slight overrepresentation of RBP-associated tripeptides in disordered regions (Fig. 3c).

The modified TriPepSVM approach predicted all 79 previously known RBPs that were detected by MS (Fig. 3d). However, 45 proteins that were previously known as RBPs according to the QuickGO annotation apparently did not shift in our gradients. It is possible that weaker and transient interactions were not preserved during fractionation of the complexes, but the presence of tRNA-interacting proteins pointed in another direction. A comparison between sedimentation velocity and molecular weight of previously known complexes showed that the gradients had a resolution limit of 50 KDa (Supplementary Fig. 5a-b), which could hamper the identification of shifting proteins that were bound to small RNAs. This limitation in the capacity for small shifts can be observed for the protein subunit of the signal recognition particle (Supplementary Fig. 5c). This protein is a well-known RBP; it has a small shift in our gradients, but the shift was not significant, according to our strict criteria. Therefore, there are a certain number of *bona fide* RBPs that were correctly predicted by TriPepSVM but not detected by R-DeeP.

On the other hand, 282 proteins were shifting in the R-DeeP analysis, but were not predicted as RBPs (Fig. 3d, Supplementary Data 7). These were RNA-associated proteins that did not bind RNA directly but were part of an RNA-binding complex, or have RNA-binding features that were not recognized by the TriPepSVM algorithm. Another 98 proteins were predicted as new RBPs, but they did not show a significant shift, and 17 proteins were predicted as new RBPs and showed a significant shift. The last group was the most promising for the identification of new RBPs. Within this group we found a DNA helicase (RuvB), ribosome factors not labeled as “RNA-binding” in UNIPROT, and also 6 proteins with no apparent previous link to RNA metabolism (All1235, All2091, Vipp1, Alr0806, Asl3888 and PrpA). To filter the data even more, we added a phylogenetic perspective to the RBP prediction. We searched for homologs of each *Nostoc* protein in 55 cyanobacterial genomes in the IMG database (see Supplementary Table S2). Next, we used our modified TriPepSVM approach to predict the conservation of the RNA-binding capacity of a protein and its homologs (Fig. 3e-f). From the subgroup of shifting proteins with high SVM scores, more than half of the Alr2890, Vipp1, Asl3888 and PrpA homologs were predicted as RBPs also in the other cyanobacteria (Fig. 3e). On the contrary, most of the homologs of Alr1700 (PatR), Alr2809, PatU3, and TrpB, proteins with a significant shift, had low SVM scores (Fig. 3e). Additionally, differences between the cyanobacterial clades could be observed in some cases. For example, Asl3888 homologs in heterocyst-forming cyanobacteria tended to have a higher SVM score or some proteins, such as PrpA, were only conserved in heterocyst-forming cyanobacteria (Fig. 3f). The complete list of SVM predictions is provided in Supplementary Data 8 and is included in our database at https://sunshine.biologie.uni-freiburg.de/R-DeeP-Nostoc/

### *In vivo* validation of six new RBPs

We used PNK assays^33^ to test *in vivo* the RNA-binding capacity of proteins from two subgroups shown in Fig. 3d; proteins shifting with high SVM scores (putative new RBPs) and proteins shifting but with low SVM scores (RNA-associated proteins). In this method, filaments containing vegetative cells and heterocysts were UV-treated to covalently crosslink the RBPs to their associated RNA(s), complexes were purified under stringent conditions to disrupt protein-protein interactions, RNA was partially digested to remove the parts of the RNA not covalently bound to the protein, and the remaining segment of the RNA was ^32^P-labeled. The RNA-protein complexes were then fractionated by SDS-PAA gel electrophoresis and transferred to a nitrocellulose membrane for autoradiography and Western Blot analysis (Fig. 4a). If a radioactive signal was detected at the position of the analyzed protein in the cross-linked sample, the protein-RNA interaction *in vivo* was confirmed.

**Fig. 4.**
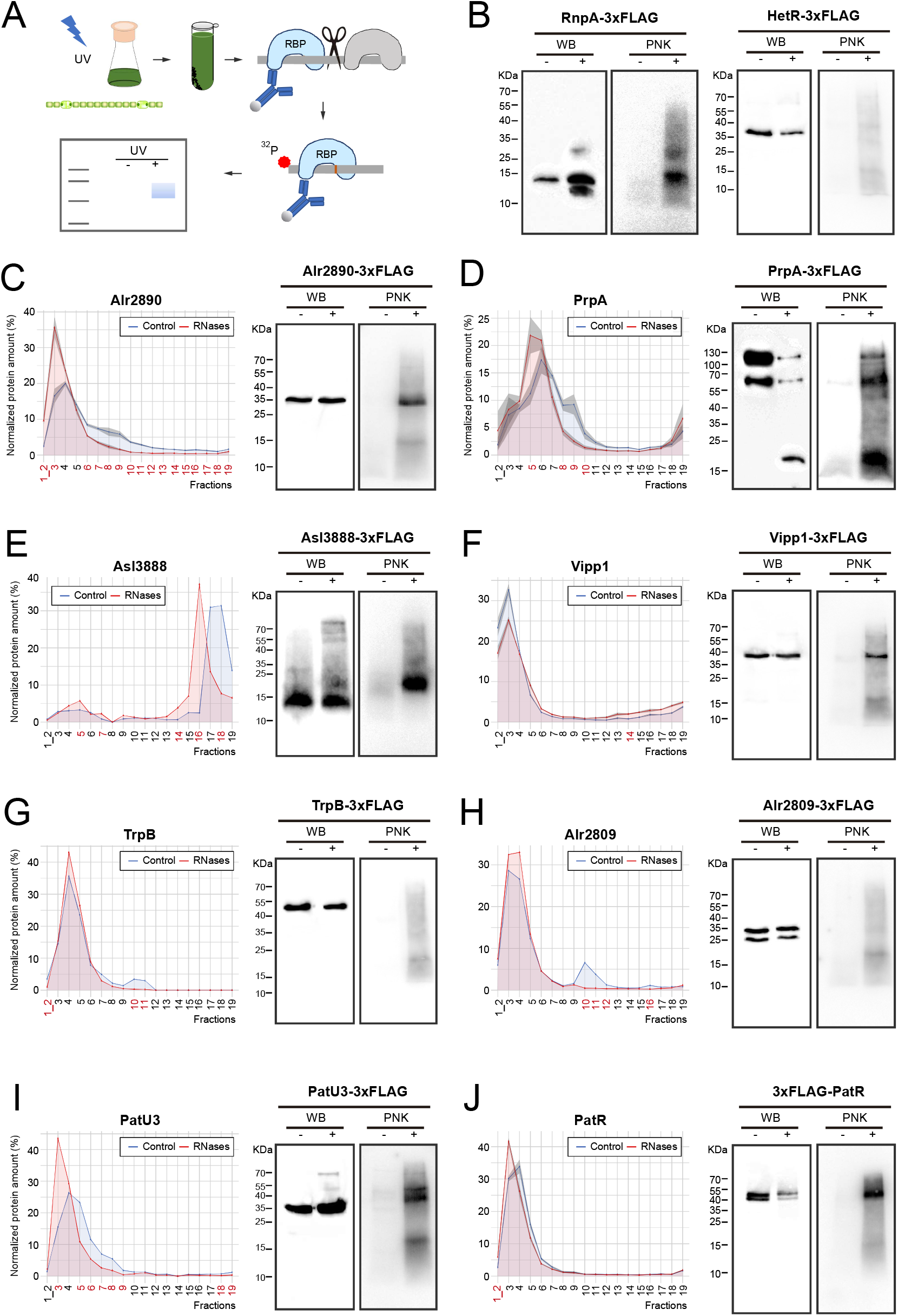
Validation of the R-DeeP screen. **a** Scheme of the PNK assays used to validate the candidate proteins. *Nostoc* filaments growing for 24h in media without combined nitrogen and containing vegetative cells and heterocysts are exposed to UV light. Filaments are lysed and proteins of interest are purified using anti-FLAG antibodies coupled to magnetic beads. Samples are treated with Benzonase to wash out other proteins that may interact with the same target RNA. The remaining RNA fragment cross-linked to the protein of interest is radioactively labelled and the protein-RNA complex is fractionated on a 15% SDS-PAA gel. The complexes are transferred to a membrane, which is later screened for radioactivity or hybridized with anti-FLAG antibody for Western blot. Non-crosslinked samples are used as controls. **b** PNK assay of RnpA-3xFLAG (positive control) and HetR-3xFLAG (negative control). It is shown the Western blot analysis of the membranes (WB) and the autoradiography (CLIP). The position of size markers (in kilodaltons) is indicated on the left side. For each candidate (**c-j**), the in-gradient distribution measured in R-DeeP (left panels) and the results of the PNK assays (right panels) are shown. The graph shows the normalized protein abundance measured by MS along the 18 analyzed fractions. The sum of protein abundance along the gradient is normalized to 100%. Raw data (mean of 3 replicates where possible) are indicated by the lines. Shading indicates the standard deviation between replicates (only if the protein was detected in more than 1 pair of untreated-treated gradients). The PNK assay panels show Western blot analysis (WB) of the membranes and autoradiography (PNK). All the Western Blot analyses were carried out using anti-FLAG antibody and the position of size markers (in kilodaltons) is indicated on the left side.

We generated *Nostoc* reporter strains that expressed each candidate protein fused to 3xFLAG epitope tags under their native promoters at their chromosomal loci, except for Asl3888 and Alr1700 (PatR) that were expressed from their native promoters but inserted in the alpha plasmid. Because photosynthetic organisms contain pigments that absorb light in the UV range, we first had to optimize the UV treatment for *Nostoc* cells (1.6 mJ/cm^2^ was finally used). As proof of concept, RnpA (an RBP) and HetR (a DNA-binding transcription factor) were used to confirm the specificity of our protocol (Fig. 4b).

The four selected proteins significantly shifting in the gradients and with high SVM scores (Alr2890, PrpA, Asl3888 and Vipp1) showed a clear radioactive signal that overlapped with the protein bands detected by Western blot on the same membrane (Fig. 4c-f), confirming their RNA-binding capacity. However, a more surprising result was obtained for PatU3 and Alr1700 (PatR) (Fig. 4 i-j), two proteins that had a shift in our gradients, but were not predicted as RBPs in any of the analyzed genomes (Fig. 3 e-f). These proteins also showed a clear radioactive signal. In contrast, two other proteins, Alr2809 and TrpB, which also had a significant shift and low SVM-scores, showed no significant labeling in the PNK assay (Fig. 4 g-h). The Western blot analysis of Asl3888 and PatU3 showed some additional faint bands of higher apparent molecular weight only in the UV cross-linked sample. A band of about 70 KDa is visible for PatU3-3xFLAG (Fig. 3i, monomer: 32 KDa) and bands between 50 and 70 KDa are visible for Asl3888 (Fig. 3, monomer: 10 KDa). These bands may indicate that these proteins may bind RNA as dimers or oligomers. The distribution of validated RBPs in the control gradients and the magnitude of their shifts after RNase treatment were also informative. For example, Alr2890 had a diffuse distribution that shifted to a sharp peak only after RNase treatment (Fig. 3c), which could be consistent with an RBP that interacts with multiple mRNAs of different lengths. In contrast, Alr1700 (PatR) had a sharp peak in the control gradients and only a very small shift after RNase treatment (Fig. 3j), which could be consistent with an interaction with only a few relatively small RNAs.

The annotated functions of some of the RBPs validated here or their homologs in other cyanobacteria are diverse and include roles in photosynthesis (Alr2890^34^, Vipp1^35^) or heterocyst differentiation (PatU3^12^, PrpA^36^), but no apparent connection to RNA has been previously described. The proteins Asl3888 and Alr1700 (PatR) lacked any functional annotation. For Asl3888 we observed an association with the 30S ribosome subunits in our cluster analysis of the control gradients (see Supplementary Data 3), but no function or association could be predicted for Alr1700 (PatR).

### PatR (Alr1700) is an RBP involved in heterocyst differentiation

Due to its enigmatic status, PatR was chosen for a more detailed analysis. According to previous transcriptomic data^37, 38^ *patR* becomes slightly downregulated after removal of combined nitrogen (Fig. 5a). These results were extended here to the protein level by observing a decrease in the PatR-3xFLAG signal after induction of heterocyst differentiation (Fig. 5b). To test this downregulation *in vivo,* we generated a reporter strain in which the promoter sequence (2038795f-2039043f of the *Nostoc* chromosome) around the transcription start site of *patR* (position 2038997f) was fused with the reporter gene *gfpmut2*. The expression of the reporter was unpatterned when the filaments were growing on plates with nitrate as a nitrogen source, while an overall significant lower fluorescence was measured at the single cell level when the filaments were growing in media without a combined nitrogen source (Fig. 5c). This result was consistent with previous transcriptomic data and our Western blot analysis (Fig. 5a-b). However, the quantification clearly showed a subpopulation of cells in which fluorescence did not decrease (Fig. 5c).

**Fig. 5.**
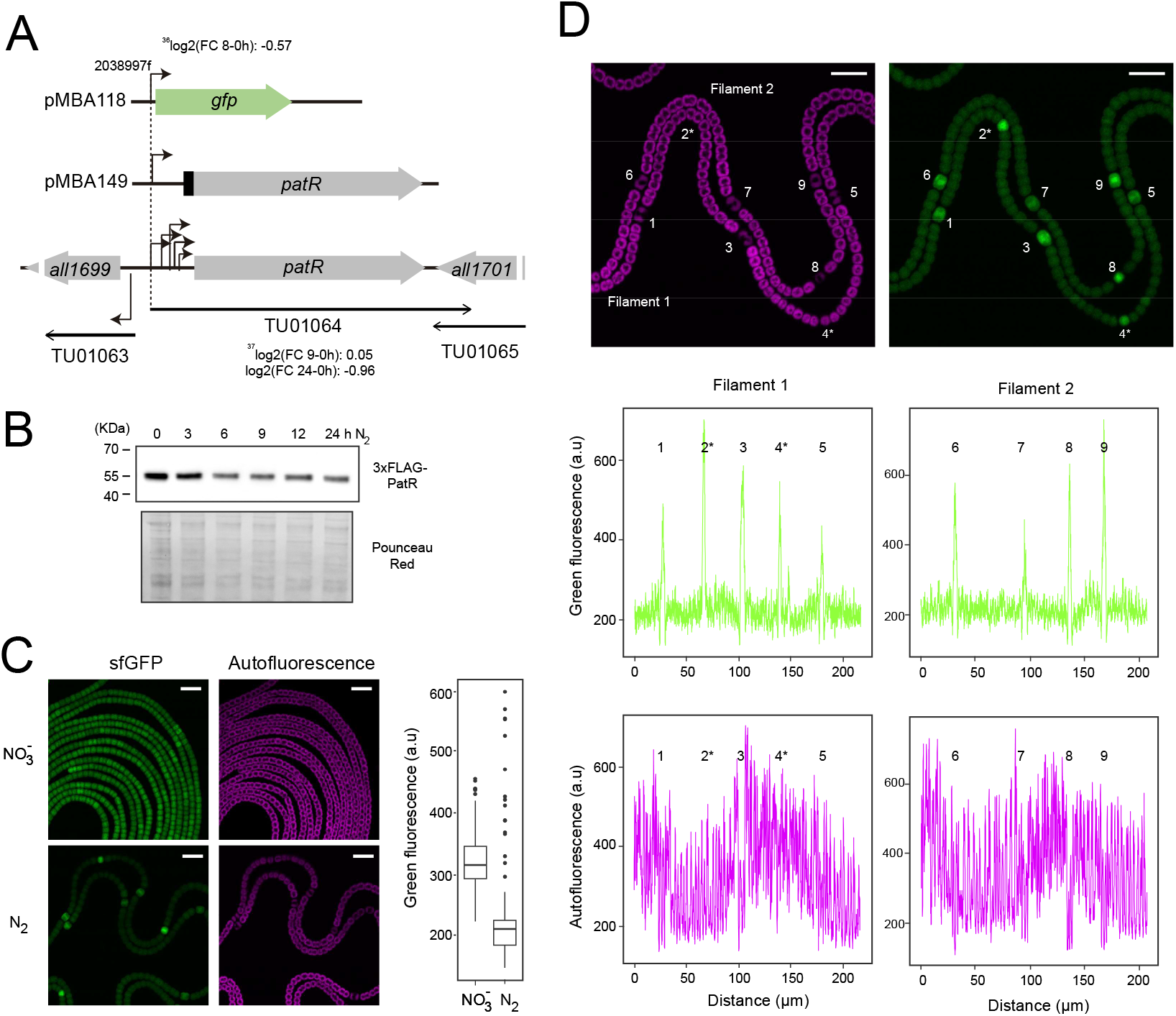
*patR* expression. **a** Scheme of DNA fragments cloned into the plasmids used to generate *Nostoc* strains with the *patR* promoter fused to *gfpmut2* (pMBA118) or 3xFLAG-PatR expressed under the native *patR* promoter (pMBA149). The diagram is not drawn to scale. Transcriptomic information from induction after nitrogen depletion is also included. Transcription start sites^37^ are indicated by bent arrows. Long lines ending in arrows indicate transcription units (TUs)^38^. **b** 3xFLAG-PatR accumulation at different time points (indicated in hours) after combined nitrogen removal. 30 µg of total soluble protein was separated on a 15% SDS-PAA gel and transferred to a nitrocellulose membrane. Anti-FLAG antibody was used. Ponceau red staining is shown as a loading control. The position of size markers (in kilodalton) is indicated on the left side. **c** Confocal fluorescence images of filaments bearing plasmid pMBA118 (*patR* promoter fusion) growing on top of medium with nitrate (NO_3_^-^) or no combined nitrogen source (N_2_). The green channel (GFP fluorescence) and the magenta channel (autofluorescence) are shown. Scale bars, 10 µm. The green fluorescence of individual cells in the two nitrogen regimes was quantified using Fiji and is shown in the adjacent boxplot (452 cells from NO_3_^-^ and 284 cells from N_2_, two-sided Wilcoxon text p-value < 2.2x10^-16^. **d** Confocal fluorescence images of filaments bearing plasmid pMBA118 growing on top of nitrogen-free medium are shown for the green channel (GFP fluorescence) and magenta channel (autofluorescence). The loss of autofluorescence from photosynthetic pigments is a characteristic feature of heterocyst differentiation. Green fluorescence and autofluorescence were quantified along two filaments using Fiji. Heterocysts and cells undergoing the heterocyst differentiation process are numbered. Cells in a very early stage of differentiation are indicated by asterisks. Scale bars, 10 µm. a.u., arbitrary units.

We analyzed in more detail the expression of *patR* in filaments growing on plates without a combined nitrogen source, a condition in which heterocyst differentiation takes place. As mature heterocysts do not perform oxygenic photosynthesis, their light-harvesting complexes are dismantled. As a result, differentiation progression can be followed at the single cell level through the reduction of chlorophyll-derived red fluorescence. Quantification along the filament of the green fluorescence of GFP and the declining red autofluorescence of developing heterocysts allows precise measurement of the timing of the cell differentiation process *in vivo^5^.* This analysis showed that the subpopulation of cells with higher fluorescence were, in fact, patterned heterocysts (Fig. 5d). Furthermore, cells at a very early stage of the development process with no emerged morphological sign (marked with an asterisk in Fig. 5d) also had a higher fluorescence than neighbouring vegetative cells. These results show that *patR* is differentially expressed between the two cell types after removal of combined nitrogen; in vegetative cells, *patR* expression is repressed, while it remains active in cells undergoing the cell differentiation process.

Due to the differential expression of *patR* between vegetative cells and differentiating heterocysts, we generated a Δ*patR* strain (Supplementary Fig. 6) to test a possible role of PatR in heterocyst differentiation. The Δ*patR* strain was unable to grow in media without combined nitrogen and this phenotype was due to the deletion since the complementation of Δ *patR* with a copy of *patR* under the *rnpB* promoter restored the growth (Fig. 6a and b lower panel). Intriguingly, the inability to grow diazotrophically was not due to a lack of heterocysts, as 24 h after combined nitrogen removal, heterocyst-like cells stained with alcian blue could be easily detected (Fig. 6b).

**Fig. 6.**
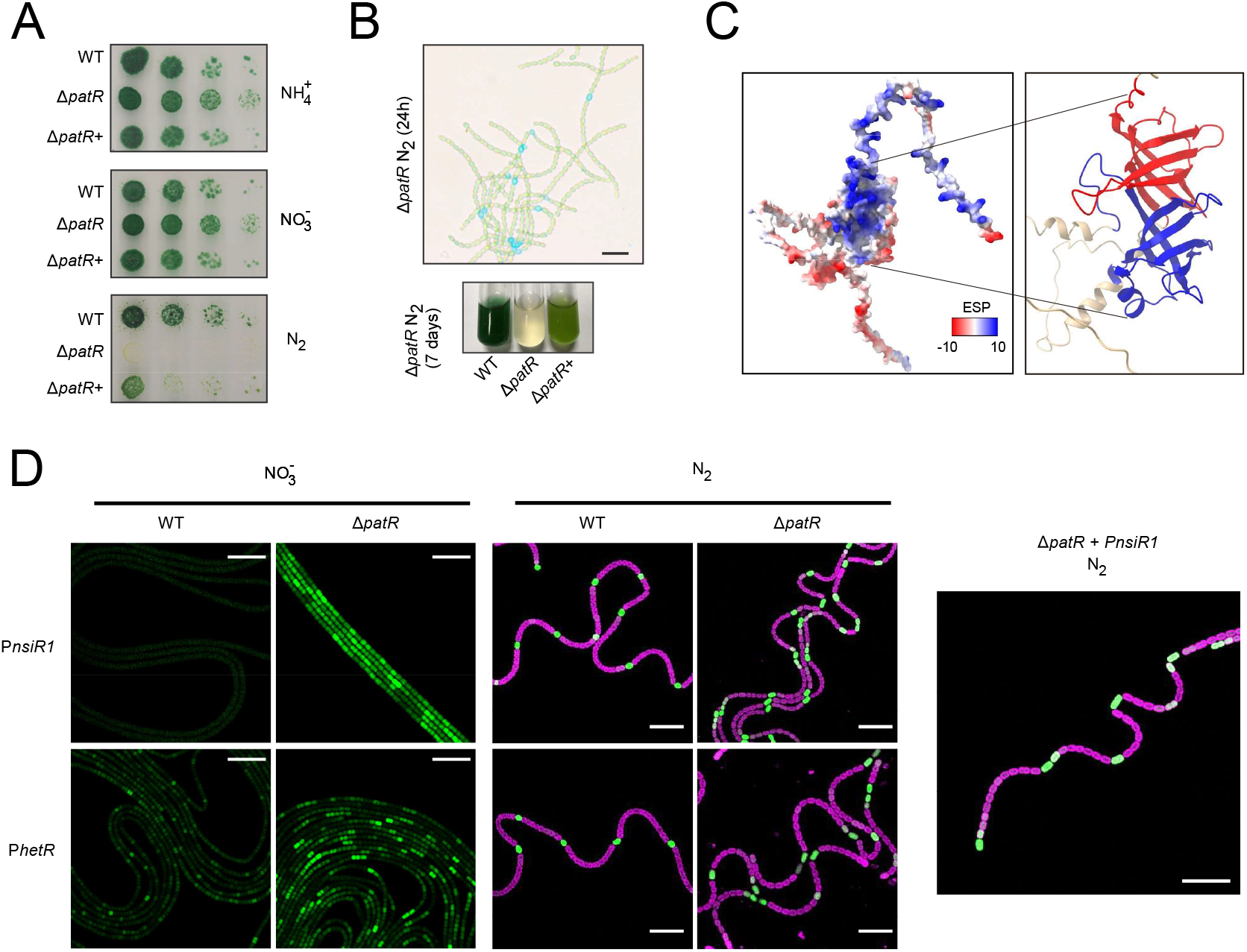
PatR regulates heterocyst differentiation. **a** Growth on solid media. Cells from liquid cultures of the indicated strains grown in the presence of nitrate were collected and resuspended in BG11_0_ at an OD_750_ = 0.1. Fivefold serial dilutions were prepared and 10 μL of each dilution plated on BG11_0_ plates containing ammonium (NH_4_^+^), nitrate (NO_3_^-^) or lacking combined nitrogen (N_2_). Pictures were taken after 13 d of incubation at 30 °C. **b** Presence of heterocyst-like cells in Δ*patR*. Upper panel: bright-field image of filaments of Δ*patR* growing in liquid medium 24h after removal of combined nitrogen. Heterocysts and heterocyst-like cells are stained with Alcian blue. Bottom panel: Growth in liquid media of indicated strains 7 d after combined nitrogen removal. **c** PatR structure predicted by Alphafold (AF-Q8YWB5-F1). Left panel: Coulombic electrostatic potential (ESP) of PatR structure calculated by ChimeraX. Positively charged surfaces (blue) and negatively charged surfaces (red) are shown. Right panel: core of the PatR protein structure. The two tandem conserved OB-fold like domains are highlighted; domain 1 in red (residues 70-158) and domain 2 in blue (residues 198-279). **d** Confocal fluorescence images of filaments from WT and Δ *patR* strains bearing promoter fusions to *gfp* of *nsiR1* promoter and *hetR* cell-type specific promoter. Filaments were grown on top of medium containing nitrate (NO_3_^-^) or without a combined nitrogen source (N_2_). Only the green channel (GFP fluorescence) is shown for the filaments growing on top of medium with nitrate. A composite image of the green channel (GFP fluorescence) and the magenta channel (autofluorescence) is shown for the filaments growing on top of medium without a combined nitrogen source (N_2_). A higher magnification image is shown on the left. Scale bars, 25 µm.

The possibility of an RBP regulating a cell differentiation process in prokaryotes was intriguing. For this reason, we decided to dissect the heterocyst differentiation process in the *patR* knockout strain in more detail. We conjugated reporter plasmids that contained the promoter of *nsiR1^39^* or *hetR^5^* fused to *gfpmut2* into Δ*patR* or WT (control). These promoters are good markers of cell differentiation, since associated genes are specifically expressed in cells at the earliest discernible stages of the differentiation process. The results indicated that the cell differentiation process in Δ*patR* was completely deregulated (Fig. 6d). In fact, the filaments of the strain Δ*patR* contained more heterocysts than the WT strain, and the increase in the number of heterocyst-like cells was so high that even patterning was affected. For example, Δ *patR* had doublets or triplets of heterocyst-like cells next to each other (Fig. 6d). This phenotype is known as Mch (multiple contiguos heterocysts) and has previously been described for mutants overexpressing HetR^40^, the master regulator of the differentiation process, or knock-out mutants in PatS^41^ or HetN^42^, negative regulators of pattern generation. Furthermore, the *nsiR1* promoter had a strong expression even if the strain was grown on plates with nitrate as the combined nitrogen source, a condition in which this promoter should not be expressed.

As a factor relevant for the regulation of heterocyst development, homologs should be expected in other cyanobacteria. Indeed, we detected *patR* homologs in 1,655 cyanobacterial genomes and 3 homologs in the endosymbiont chromatophore genomes of photosynthetic *Paulinella^43^*. The alignment of 107 selected homolog sequences from a wide diversity of cyanobacteria revealed two conserved domains (residues 70-158 and 198-276 of PatR, see Supplementary Data 9). The PatR structure predicted by Alphafold has to β barrels resebling OB-fold domains that correspond to the two conserved domains (Fig. 6c). A fold-seek^44^ search revealed that one of the domains was structurally related to OB-fold containing proteins that bind RNA, while the second domain was more similar to OB-fold containing proteins that bind ssDNA. This search is consistent with our data and with the fact that a prediction of the electrostatic potential of the PatR structure revealed a highly positive potential on one of the faces of the two beta barrels (Fig. 6c), indicating the potential RNA binding site.

The role of PatR in heterocyst differentiation initially appeared to be inconsistent with its universal conservation in all types of cyanobacteria. However, phylogenetic analysis revealed that homologs in heterocyst-forming cyanobacteria form a coherent clade clearly distinct from the other cyanobacteria (Supplementary Fig. 7). Indeed, structural comparison between PatR and its homologs in two unicellular species, *Synechocystis* sp. PCC 6803 (hereafter *Synechocystis*) and *Prochlorococcus marinus* MED4 revealed that the separation of homologs in heterocyst-forming cyanobacteria was based on the presence of two additional α-helices (Supplementary Fig. 8).

These results establish that a widely conserved RBP with two OB-fold domains is involved in the regulation of heterocyst differentiation.

## Discussion

Here, we have applied an R-DeeP/TripepSVM hybrid approach to find RBPs in a prokaryote undergoing a cell differentiation process. The most crucial point for the correct performance of this technique is the preservation of the protein-protein and RNA-protein complexes in the untreated samples, while the RNA should be digested completely in the RNase-treated samples. Under the conditions tested, RNA appeared to be intact in the untreated gradients (Fig. 1c), and the distribution of highly expressed RNA species was similar to that found in a Grad-seq dataset in the unicellular cyanobacterium *Synechocystis*^45^. Furthermore, the overall similar protein profiles of the untreated and treated gradients visualized by SDS-PAA gel electrophoresis (Supplemental Fig. 1c-d) plus the detection of a clear shift for the ribosomal protein RplA, but not for the DNA-binding HetR confirmed the specificity of our approach (Fig. 1e).

Based on the R-DeeP data alone, 333 RNA-associated proteins were identified (Fig. 3), which constitutes 12% of the detected proteome. A related approach, GradR, identified 3% of the detected proteome as RNA-associated in *S. typhimurium*^21^, whereas even 37% RNA-associated proteins were detected in human cells^46^. Alternative approaches detected 364 and 384 new RBPs in *E. coli*^47^ and *Staphylococcus aureus*^48^, respectively. Therefore, the number of RNA-associated proteins identified by our approach is in a range similar to that of other prokaryotic proteomes.

Half of the identified RNA-associated proteome moved to lower fractions (lower molecular weight) after RNase treatment, while the other half moved to higher fractions (higher molecular weight) (Fig. 3a). Although this behavior could seem striking, it has also been observed in the GradR/R-DeeP analyses of *Salmonella* or human samples^21, 46^. One possible explanation is the precipitation of a protein when its RNA counterpart is not present. In the heterologous purification of some RBPs, such as CRISPR proteins, it was observed that the coexpression of their target RNA was required to stabilize the proteins in soluble form ^49^. However, the right-shift could also reflect a regulatory mechanism (see the discussion below about Vipp1).

The prediction of RBPs usually relies on the search for conserved RNA-binding domains. However, the ability to bind RNA may be based on subtler patterns. Therefore, we used a modified version of TriPepSVM^32^, an algorithm that is trained with known RBPs or non-RBPs sequences, according to the databases, and considers their tripeptide content for RBP prediction. The advantage of this method is the ability to predict new RBPs whose RNA binding sites could be located in intrinsically disordered regions. In fact, we observed a slight tendency of the characteristic RBP tripeptides to be located in disordered regions in *Nostoc* (Fig. 3c). This tendency was previously reported for the human proteome, but not for the proteomes of *E. coli* and *S. typhimurium*^32^. The importance of disordered regions in RBPs is a recent field of research^50^ since some RBPs are assembled in liquid-liquid phase separation droplets^51^, including an RNA helicase in cyanobacteria^52^.

There was only a partial overlap between the shifting proteins and the prediction of TriPepSVM (Fig. 3d), indicating complementarity between the two methods. The shifting proteins not predicted as RBP could be part of RNA-protein complexes but may not bind RNA themselves. The presence of previously known RBP with no apparent significant shift has also been reported^21^. Weak and transient RNA-protein interactions or the R-DeeP resolution limits (Supplementary Fig. S5) could be the reasons for these false negatives. Therefore, some of the proteins not significantly shifting but predicted as RBPs might be true RBPs. Two metabolic enzymes related to nitrogen metabolism, CphA1, cyanophycin synthetase^53^, and UreB, an urease subunit^54^, had a visible slight shift in our gradients just below our strict cutoff, but in contrast had very good SVM scores not only in *Nostoc* but in all cyanobacterial proteins tested (Fig. 3e-f). They could be false positives predicted by TriPepSVM, but a recent report found both proteins as interacting partners of YlxR, an RBP in *Synechocystis*^55^. The detection of new RNA binding capacities in many metabolic enzymes has also been reported in *S. typhimurium*^48^, and may represent a direct link between responses to nutrient availability and post-transcriptional gene regulation.

All four proteins with significant shifts and high SVM scores (Alr2890, Vipp1, PrpA, and Asl3888) had the ability to interact with RNA *in vivo* (Fig. 4c-e), confirming the specificity of our combined approach. However, we also obtained very clearly labeled bands for PatU3 and PatR (Fig. 4i-j), two proteins with low SVM scores in most of the cyanobacterial proteomes. This result suggests that some of the 282 proteins that shifted but were not predicted to be RBPs (Fig. 3d) could be true RBPs. It is important to note that SVMs are trained with current knowledge and, therefore, totally new domains or interaction patterns could be missed.

The newly validated RBPs have previously annotated functions related to essential physiological processes such as photosynthesis and heterocyst differentiation. The Alr2890 homologs are strictly conserved in cyanobacteria and plant chloroplasts. Although Alr2890 contains a putative S4 RNA binding-domain and was therefore tagged as “RNA-binding” in the UNIPROT database, Sll1252, its close homolog in *Synechocystis*, was found in PSII preparations, is necessary for the proper activity of PSII^34^ and is involved in the coordination of electron transport between plastoquinone and the cytochrome b_6_f complex^56^. The authors hypothesized a regulatory function for Sll1252^34^, but the exact mechanism has remained unknown. Here, we confirm that the *Nostoc* ortholog Alr2890 is an RBP, suggesting that it is bifunctional.

The case of Vipp1 (All2342) is even more striking, since it was the only validated RBP that had a shift toward higher fractions after RNase treatment (Fig. 4f). Oligomers of Vipp1 are essential for maintaining the integrity of the thylakoid membrane^35^ and Vipp1 was recognized as a bacterial ancestor of the eukaryotic ESCRT-III membrane remodeling system^57^. In cyanobacteria, the protein continuously moves between a diffused fraction and discrete foci at the thylakoid membranes^58^, which are more abundant after high light stress^59^ and consist of Vipp1 oligomers^35^. The trigger that causes the polymerization of the protein is unknown. Based on our observation that Vipp1 migrated in our gradients after RNase treatment to multiple fractions corresponding to higher molecular weight, it is tempting to speculate that it is the presence of an RNA that sequesters Vipp1 in monomeric form. After the removal of this hypothetical RNA (or degradation in our treatment), Vipp1 is able to form oligomers.

Proteins involved in heterocyst differentiation were also validated as new RBPs. PrpA was first described as a phosphatase that affects heterocyst differentiation^36^. Here, we have demonstrated the *in vivo* RNA-binding capacity of PrpA, which fits the report that a closely related homolog in *Trichormus variabilis* ATCC29413 is part of an RNA repair system^60^. In contrast, no link to RNA has been reported for PatU3, a protein expressed specifically in heterocysts^61^ that is involved in the regulation of heterocyst formation and cell division^12^.

Our mutation of *patR* highlighted the fundamental role that this RBP plays in the heterocyst differentiation process (Fig. 6). The absence of *patR* caused a total deregulation of heterocyst patterning and differentiation (Fig. 6). Δ*patR* filaments had a higher tendency to differentiate heterocysts, since even growing in plates with nitrate, a condition in which the differentiation process should not occur, the strain had a higher expression of *nsiR1* and *hetR* (Fig. 6d). After combined nitrogen removal, this tendency for cell differentiation generated an unnatural high number of heterocyst-like cells (Fig. 6d), sometimes control of the patterning is lost and generates doublets and triplets of heterocyst-like cells. It is intriguing that this phenotype of multiple contiguous heterocysts (Mch) was reported for cells lacking PatU3^61^, also characterized as RBP here. This higher frequency of heterocyst-like cells did not improve diazotrophic growth, but rather resulted in an inability to grow diazotrophically at all. We consider two explanations for this observation. Heterocysts may not have been able to complete maturation; alternatively, the ratio of heterocysts to vegetative cells may not have been appropriate for nitrogen assimilation, since nitrogen is assimilated via the GS-GOGAT cycle and sufficient amounts of carbon skeletons must be provided by neighboring vegetative cells.

These strong phenotypic effects suggest PatR as a key regulator of heterocyst maturation. Interestingly, there is also a parallel to HetR, the transcriptional master regulator of heterocyst differentiation. The *hetR* gene is transcribed from four discernible start sites^62^. Apparently, *patR* expression decreased after combined nitrogen removal (Fig. 5a-b), but our fluorescence confocal analysis of single cells revealed a more complex picture (Fig. 5c). Expression of this gene appeared to be repressed from early stages of heterocyst differentiation only in cells that were not undergoing the cell differentiation process (Fig. 5d). Although we analyzed the transcription from the main transcriptional start site of *patR* (Fig. 5a), four more transcriptional start sites were identified for this gene^37^. The presence of multiple transcriptional start sites, sometimes with paradoxical regulations that create complex expression patterns along the filament, has also been observed for other differentiation-relevant genes in *Nostoc*^63^. Therefore, the complex promoter architecture is an intriguing parallel between PatR and other genes critical for the control of heterocyst differentiation.

The wide phylogenetic conservation of PatR suggests that it is also important for non-heterocystous cyanobacteria. The protein has a structural architecture that contains two OB-fold domains combined in tandem (Fig. 6c). A similar arrangement has been described for an ssDNA binding protein in *Deinococcus radiodurans*^64^. Although the presence of PatR homologs in unicellular cyanobacteria could be paradoxical considering its effects on heterocyst differentiation, homologs of other proteins essential for heterocyst differentiation, such as HetR, also exist in non-heterocyst-forming cyanobacteria^65^. PatR and its homologs probably have some essential function related to the physiology of all cyanobacteria, but the insertion of to additional α-helices in the homologs of heterocyst-forming cyanobacteria (Supplementary Figures 7 and 8) may have allowed the particular regulatory role of PatR in heterocyst differentiation. The exact regulatory mechanism(s) involving PatR and its precise RNA targets are highly interesting topics for future work.

## Methods

### Strains and growth conditions

The different *Nostoc* sp. PCC 7120 strains used in this work (Supplementary Table S3) were grown photoautotrophically with constant shaking under standard conditions (30 °C and 50 µE illumination) in liquid BG11 medium^66^. Filaments were collected by centrifugation at 3,270 x g for 5 min at room temperature, washed and then resuspended in BG11_0_ (medium without a combined nitrogen source) to induce heterocyst differentiation. *Nostoc* sp. PCC 7120 derivative strains bearing Sm^R^Sp^R^ or Nm^R^ plasmids were grown in liquid in the presence of 2 µg/mL streptomycin (Sm) and spectinomycin (Sp) each, or 5 µg/mL neomycin (Nm).

### R-DeeP experiment; cell lysis, gradient preparation and fractionation

Triplicate 200 mL *Nostoc* liquid cultures were grown photoautotrophically under standard conditions in BG11 medium until they reached a concentration of 3 µg chlorophyll /mL. Heterocyst differentiation was induced as described above. After 26 h of cultivation, the filaments were harvested by centrifugation and resuspended in ice-cold lysis buffer (20 mM Tris-HCl pH 7.5, 150 mM KCl, 10 mM MgCl_2_, 1mM DTT) supplemented with protease Inhibitor (cOmplete EDTA-free, Roche). From this point on, the samples were kept at 4 °C. Each replicate was divided into equal sample tubes. The control samples in which the RNA should be preserved were supplemented with 8 µL of Ribolock RNase Inhibitor. To all tubes, 100 µL of a mixture of 0.1 mm and 2.5 mm glass beads were added. Cells were mechanically lysed in a pre-chilled Precellys homogenizer (Bertin Technologies) using 9 cycles of 10 s shaking at 6,000 rpm followed by a pause of 5 s. Cell debris and glass beads ere removed by centrifugation at 1,500 x g for 2 min. Next, 1.25% β-DM was added to the supernatant to solubilize membrane proteins by vortexing for 30 s. Membranes were pelleted by centrifugation at 21,000 x g for 15 min. Next, 100 µL of RNase A/T1 mix (Thermo, 2 mg/mL RNase A and 5,000 U/mL RNase T1 stock) was added to the RNase-treated extracts, while 100 µL of RNase buffer (50 mM Tris-HCl pH 7.4 and 50% glycerol) was added to the control extracts. Control and RNase-treated samples were incubated on ice for 20 min.

Linear sucrose gradients were prepared on Ultra-Clear Beckman tubes using a Gradient Master 108 forming system (Biocomp) and two solutions of lysis buffer supplemented with either 10% or 40% sucrose. 600 µL of the control and RNase-treated total extracts were loaded carefully onto the gradients. Macromolecular complexes were separated by ultracentrifugation of the lysates in a swinging-bucket rotor (Beckman SV40 Ti) for 16 h at 285,000 x g. Fractions of equal volume (600 µL) were collected using a Piston Gradient Fractionator (Biocomp). The protein concentration of each fraction was determined using the Lowry procedure^67^.

### Northern and Western blot analysis of fractions

RNA was extracted using hot phenol^68^ with modifications^6^ from 300 µL of each fraction. The RNA was fractionated on a 10% urea-PAA gel, stained with ethidium bromide, and electroblotted onto a Hybond-N+ membrane (Amersham) at 1 mA per cm^2^ for 1 h. The distribution of RnpB, Yfr1 and NsiR8 was analyzed by Northern hybridization using single-stranded radioactively labeled RNA probes transcribed *in vitro* from PCR-generated templates (see Supplementary Table 2 for primers). Radioactively labeled probes were generated using [α -^32^P]-UTP and the Maxiscript T7 In vitro transcription kit (Thermo Fisher Scientific). The distribution of NsiR1, 4.5S and 5S along the gradient fractions was also analyzed by Northern hybridization using end-labeled oligonucleotides (see Supplementary Table 2 for primers). Oligonucleotides were labeled iw th [γ -^32^P]-ATP and polynucleotide kinase (PNK). Thermo). Hybridizations were performed as previously described^69^. The signals were detected with a Typhoon FLA 9500 (GE Healthcare).

For Western blot analysis, 35 µL of each fraction were fractionated in 15% SDS-PAA gels and transferred to a nitrocellulose membrane (Amersham). As a control, identically prepared gels were stained with Coomassie Brilliant Blue G-250. The membranes were blocked in 5% skim milk in T-PBS (1% Tween in PBS) for 1 h at room temperature and incubated with rabbit polyclonal antibody (#AS111738,1:5000, Agrisera) or mouse monoclonal anti-FLAG coupled to HRP (#A8592,1:2000, Sigma) for 16 h at 4 °C. The membranes were washed three times with T-PBS, 5 min each. Membranes previously incubated with anti-RplA were incubated with the secondary antibody anti-rabbit IgG-HRP (#A8275, 1:10000, Sigma) for 1 h at room temperature and then washed three times with T-PBS. Signals were detected with Western-Blot ECL spray (Advanta).

### Sample preparation for MS and measurements

Samples for proteomic analysis were prepared as previously described^70^, with minor modifications. 1% N-lauroylsarcosine was added to equal volumes of samples along the gradient (fractions 1 and 2 were combined) and proteins were digested following a modified SP3 protocol^71^. Specifically, 20 µL samples were incubated with 40 µL bead suspension (Thermo Fischer Scientific, Sera-Mag Speed Beads, 4515-2105-050250, 6515-2105-050250) in 2.5% formic acid and 50% ethanol (15 min, RT, 500 rpm). Beads were then washed 4x with 70% ethanol and proteins were digested overnight (RT, 500 rpm) with trypsin and LysC (200 ng of each enzyme in 5 mM chloroacetamide, 1.25 mM TCEP in 100 mM HEPES pH 8 per 5 ug protein). Peptides were eluted from the beads, dried under vacuum, and resuspended in water. Ten microliter samples (up to 5 µg per sample) were labeled with 50 µg of TMT18plex (Thermo Fisher Scientific) reconstituted in 4 µL acetonitrile. Labeling reaction (1h, RT, 500 rpm) was quenched with 4 µL 5% hydroxylamine (30 min, RT, 500 rpm) and samples of the same gradient were pooled. The samples were then desalted with solid phase extraction by loading onto a Waters OASIS HLB μElution Plate (30 μm), washing twice with 100 μl of 0.05% formic acid, and eluting with 100 μl of 80% acetonitrile. Samples were then dried under vacuum and resuspended in 20 mM ammonium formate pH 10.0.

Offline high pH reverse phase fractionation was performed using an Agilent 1200 Infinity high-performance liquid chromatography (HPLC) system equipped with a quaternary pump, degasser, variable wavelength UV detector (set to 254 nm), Peltier-cooled autosampler, and fraction collector (both set at 10 °C for all samples). The column was a Gemini C18 column (3 μm, 110 Å, 100 x 1.0 mm, Phenomenex) with a Gemini C18, 4 x 2.0 mm SecurityGuard (Phenomenex) cartridge as a guard column. The solvent system consisted of 20 mM ammonium formate (pH 10.0) (Buffer A) and 100% acetonitrile as mobile phase (Buffer B). The separation was accomplished at a mobile phase flow rate of 0.1 mL/min using the following linear gradient: 100% Buffer A for 2 min, from 100% Buffer A to 35% Buffer B in 59 min, to 85% Buffer B in a further 1 min, and held at 85% Buffer B for an additional 15 min, before returning to 100% Buffer A and re-equilibration for 13 min. Forty-eight fractions were collected along with the LC separation and subsequently pooled into 12 fractions. Pooled fractions were dried under vacuum centrifugation, reconstituted in 10 μL 1% formic acid, 4% acetonitrile prior LC-MS analysis.

An UltiMate 3000 RSLC nano LC system (Dionex) fitted with a trapping cartridge (µ-Precolumn C18 PepMap 100, 5µm, 300 µm i.d. x 5 mm, 100 Å) and an analytical column (nanoEase™ M/Z HSS T3 column 75 μm x 250 mm C18, 1.8 μm, 100 Å, Waters) was used. Trapping was carried out with a constant flow of 0.05% trifluoroacetic acid at 30 µL/min onto the trapping column for 6 minutes. Subsequently, peptides were eluted via the analytical column with a constant flow of solvent A (3% DMSO, 0.1% formic acid in water) at 0.3 µL/min with an increasing percentage of solvent B (3%DMSO, 0.1% formic acid in acetonitrile). The outlet of the analytical column was coupled directly to a QExactive plus (Thermo) mass spectrometer using the anospray Flex™ ion source in positive ion mode.

The peptides were introduced into the QExactive plus via a Pico-Tip Emitter 360 µm OD x 20 µm ID; 10 µm tip (CoAnn Technologies) and an applied spray voltage of 2.3 kV. The capillary temperature was set at 320°C. Full mass scan was acquired with a mass range of 375-1,200 m/z in profile mode with resolution of 70,000. The filling time was set at a maximum of 250 ms with a limitation of 3x106 ions. Data dependent acquisition (DDA) was performed with the resolution of the Orbitrap set to 35,000, with a fill time of 120 ms and a limitation of 2x105 ions. A normalized collision energy of 30 was applied. The isolation window of the quadrupole was set to 0.7 m/z. A dynamic exclusion time of 30 s as used. The peptide match algorithm was set to ‘preferred’ and charge exclusion ‘unassigned’, charge states 1, 5 - 8 were excluded. MS2 data was acquired in profile mode.

### Bioinformatic analysis of R-DeeP data

Bioinformatic analysis was performed in R. Only proteins with at least two identified peptides were kept for the analysis. In the case of small proteins (< 100 aa), a more relaxed filter was applied and a single identified peptide was considered sufficient to keep the protein in the analysis. Furthermore, only proteins identified in at least one pair of control and RNase-treated gradients were kept. Batch effect removal and a variance stabilisation normalization method were applied to each fraction using the *limma^31^* and *vsn^72^* packages. An in-gradient normalisation was also performed, so that the sum of each protein along a gradient was set to 100%. A differential expression-like approach was applied to each fraction using the *limma* package. The presence of a protein in multiple replicates was taken into account using the parameter *weights* for the adjustment of the linear model through the *lmFit* function. The false discovery rate was calculated using the *fdrtools^73^* package. Proteins with a fold change ≥ 2 and fdr ≤ 0.05 in at least one fraction ere considered as hits. Proteins iw th a fold change ≥ 1.5 and fdr ≤ 0.02 ere considered candidates.

An R package with the annotation of *Nostoc* was created using the *AnnotationForge* R package. Sequences and description of the *Nostoc* proteome were downloaded from NCBI and UNIPROT. The GO terms were downloaded from QuickGO (https://www.ebi.ac.uk/QuickGO/annotations). The molecular weight and isolelectric point for each protein were calculated using the Expasy *Compute pI/Mw tool* (https://web.expasy.org/compute_pi/). GO-term enrichment of the RNA-associated proteome was performed using the *clusterProfiler*^74^ R package and a q value ≤ 0.05 was used as cutoff.

In order to group proteins with similar sedimentation profiles, a soft clustering approach was applied to the mean of the control gradients by Mfuzz^75^. A m parameter of 1.14 was selected and a confidence greater than 0.85 was used as the threshold for a protein to be considered in the core of a cluster. The log2-fold change of each fraction for the 333 RNA-associated proteins was clustered using hierarchical clustering. Silhouette was used as a metric to determine the best number of clusters in both cases.

### Bioinformatic prediction of RBPs

We used a modified version of TriPepSVM^32^ to predict RBPs in cyanobacteria. Instead of using a reference proteome and its evolutionary close proteomes for the generation of positive and negative datasets, we selected proteomes of cyanobacteria with diverse morphologies and lifestyles (see Supplementary Table S1). In addition, we also included the proteome of two well-studied Gram-negative bacteria, *E. coli* K12 and *S. typhimurium* LT2. All proteomes were downloaded from UNIPROT. The assembly of the positive and negative training datasets was performed following the TriPepSVM pipeline^32^. We used the KeBABS^76^ package to perform a 10-k-fold validation in a grid search using the following combination of parameters: positive class weight, negative class weight, k-mers, and cost. The balanced accuracy score (BACC) was used to select the best combination of parameters. An RBP prediction was performed using the following parameters: positive class weight = 2.7, negative class weight = 0.05, cost = 1 and k-mer = 3. A threshold of 0.25 was selected in the SVM scores of the proteins for classification as potential RBP, resulting in 311 potential RBPs.

The prediction of intrinsically disordered regions in the predicted RBPs was performed using IUPred2A^77^. Regions with an average disorder score greater than 0.5 were considered not globular regions. A sliding window programmed in R was used to count the frequency of tripeptides in globular or disorder regions of the proteins. A curve was smoothed using LOESS regression (span = 0.2) to analyze the tendency of tripeptides enriched in RBPs to localize in intrinsically disordered regions.

To investigate the evolutionary conservation of the RNA binding potential of *Nostoc* proteins, homologs were predicted for the entire *Nostoc* genome using “Genome Gene Best Homologs” from the IGM/MER Portal of the IMG database^78^, 55 cyanobacterial genomes (see Supplementary Table S2), a minimum percentage of identity of 40% and an e-value of 0.05. A prediction of RBPs for all homologs was carried out using the parameters specified above.

### Construction of mutant and reporter strains

To generate a Δ*patR* strain, two overlapping fragments were amplified by PCR using genomic DNA with oligonucleotides 669 and 670 and oligonucleotides 671 and 672, respectively (see Supplementary Table S4 for oligonucleotides). The resulting products were then used as templates for a third PCR with oligonucleotides 669 and 672 resulting in deletion of the sequences corresponding to *patR* and the generation of a unique XhoI site between the two amplified fragments. The fragment was digested with BamHI at the sites provided by oligonucleotides 669 and 672, and cloned into BamHI-digested *sacB*-containing Sm^R^Sp^R^ vector pCSRO^79^, yielding pMBA62 (see Table S5 for plasmids). An Nm^R^ cassette was excised from pRL278^80^ as a SalI-XhoI fragment and cloned at the Xho site of pMBA62, yielding pMBA66, which was transferred to *Nostoc* by conjugation^81^. Genomic DNA was extracted from clones exhibiting sucrose resistance and sensitivity to neomycin. Segregation of the clones was analyzed by PCR using genomic DNA and oligonucleotides 738 and 739. Clones without the *patR* gene and no wild-type chromosome ere named Δ *patR*.

The plasmid pMBA79 was constructed to express *patR* from the *rnpB* promoter. A PCR fragment was amplified using oligonucleotides 738 and 739, digested with NsiI and XhoI and cloned in NsiI and XhoI-digested pMBA20^82^. pMBA79 and two reporter plasmids (pSAM301^39^ and pSAM270^5^) were transferred to Δ*patR* strain by conjugation with selection of Sm^R^Sp^R^ cells.

Two plasmids for the expression of proteins fused with 3x-FLAG were designed (see the supplementary Table S5 for the plasmid descriptions). Fragments containing *sgfp* fused to the 3xFLAG sequence were amplified by PCR using as template pMBA96^8^ and oligonucleotides 32 and 34 (3xFLAG N-terminal fusion) or 33 and 35 (3xFLAG C-terminal fusion). The resulting products were NsiI/SacI-digested and cloned into NsiI/SacI-digested pMBA20^82^, yielding pMBA131 and pMBA132, respectively. The *rnpB* promoter and *sfGFP* of pMBA132 were exchanged by upstream sequences plus the coding sequences of the candidate RBPs. DNA fragments were amplified by PCR using as template genomic DNA and oligonucleotides 129 and 130, 136 and 137, 148 and 149, 151 and 152, 156 and 157, 166 and 167, 194 and 195 or P03 and P04, resulting in the amplification of upstream sequences plus the coding sequences of *alr2809*, *vipp1*, *patU3*, *prpA*, *trpB*, *rnpA*, *alr2890* and *asl3888*, respectively. Fragments were PstI/XhoI-digested and cloned into PstI/XhoI-digested pMBA132, yielding pMBA166, pMBA170, pMBA177, pMBA179, pMBA182, pMBA188, pMBA211 and pMB2, respectively. All plasmids contained the upstream sequences plus the coding sequences of the selected genes fused to the 3XFLAG sequence and were designed to integrate in the alpha plasmid of *Nostoc*. DNA fragments were amplified by PCR using as template pMBA166, pMBA170, pMBA177, pMBA179, pMBA182, pMBA188 or pMBA211, forward oligonucleotides 131, 138, 150, 153, 158, 168, 196 and 19 as reverse oligonucleotide, respectively. These fragments were digested with SacI and cloned into SacI-digested pCSV3^83^, yielding pMBA167, pMBA171, pMBA178, pMBA180, pMBA183, pMBA189 and pMBA213, respectively. These plasmids contain the upstream sequences plus the coding sequences of *alr2809*, *vipp1*, *patU3*, *prpA*, *trpB*, *rnpA,* and *alr2890* fused to 3XFLAG sequence and allowed the insertion of the fragments at the original chromosomal loci expressing the fusion proteins from their native promoters. Plasmids pMBA167, pMBA171, pMBA178, pMBA180, pMBA183, pMBA189, pMBA213 and pMB2 were transferred to *Nostoc* by conjugation with selection of Sm^R^Sp^R^ cells.

An N-terminal fusion of PatR to the 3xFLAG epitope was generated. A fragment containing the promoter and 5’UTR of *patR* was amplified by PCR using genomic DNA as template and oligonucleotides 57 and 58. The resulting product was digested with PstI and NsiI and cloned into PstI/NsiI-digested pMBA131, yielding pMBA140. A fragment containing the coding region of *patR* was amplified by PCR using as template genomic DNA and oligonucleotides 53 and 54. The product was digested with XhoI and SacI and cloned into XhoI-/SacI-digested pMBA140, yielding pMBA149, which was transferred to Δ*patR* by conjugation with selection of Sm^R^Sp^R^ cells.

In order to generate a partR promoter reporter fused with *gfpmut2*, a fragment containing the promoter region of *patR* (positions 2038795f to 2039043f) was amplified by PCR using genomic DNA as template and oligonucleotides 978 and 979. The resulting fragment was digested with ClaI/XhoI and cloned into ClaI/XhoI-digested pSAM270^5^, yielding pMBA117, which was transferred to *Nostoc* by conjugation with selection of Sm^R^Sp^R^ cells.

A markerless HetR-3xFLAG strain was generated. A fragment containing the promoter, 5’TR and coding region of *hetR* was amplified by PCR using as template genomic DNA and oligonucleotides 49 and 70. The resulting fragment was PstI/XhoI-digested and cloned into PstI/XhoI-digested pMBA132, yielding pMBA154. A fragment containing the sequences downstream of *hetR* was also amplified by PCR using genomic DNA as template and oligonucleotides 50 and 51. The fragment was digested with SacI at the sites provided by the oligonucleotides and cloned into SacI-digested pMBA154, yielding pMBA157 (only clones with the correct orientation of the downstream region were kept).

A fragment containing the full construct (promoter, 5’T, *hetR* coding region, 3xFLAG, and downstream region) was finally amplified by PCR using pMBA157 as template and oligonucleotides 47 and 52. The resulting fragment was digested with BamHI at the site provided by the oligonucleotides and cloned into BamHI-digested pCSRO^83^, yielding pMBA160, which was later transferred into the Δ*hetR* mutant CSSC2^84^ by conjugation. Sm^S^Sp^S^ clones exhibiting resistance to sucrose and able to grow again diazotrophically were selected and sequenced to confirm the presence of a *hetR* copy fused to the 3xFLAG inserted at the *hetR* locus.

### PNK assays

The PNK assays were carried out as previously described^33^ with some modifications. 400 mL of cultures from reporter strains expressing 3xFLAG protein fusions were grown in BG11 supplemented with the appropriate antibiotic until they they reached a concentration of 2.5 µg chlorophyll/mL. Heterocyst differentiation was induced as described above. After 24 h of cultivation, half of each culture (200 mL) was treated (1.6 mJ/cm^2^) while gently shaking to induce RNA-protein crosslinking keeping the sample on ice (the remaining half was also placed on ice). Cells were pelleted by centrifugation at 3,270 x g for 5 min. The pellets were resuspended in NT-P buffer (50 mM NaH_2_PO_4_, 300 mM NaCl, 0.05% Tween, pH 8.0) supplemented with protease Inhibitor (cOmplete EDTA-free, Roche). Cells were lysed as described for the R-DeeP experiment. Glass beads were removed by centrifugation at 1,500 x g for 2 min. Next, membranes were pelleted by centrifugation at 21,000 x g for 30 min at 4 °C. 2 mL of clarified lysate per sample (crosslinked and non-crosslinked) was incubated with 80 µL of anti-FLAG M2 magnetic beads (Sigma) prewashed 5 times in NT-P buffer. From this point on, all the washing steps were performed with 2 mL of buffer. The samples were incubated at 4 °C while gently rotating for 1 h. Beads were washed twice with high-salt NT-P buffer (50 mM NaH_2_PO_4_, 1 M NaCl, 0.05% Tween20, pH 8.0), once with of cold NT-P buffer and once with benzonase buffer (50 mM Tris-HCl, 1 mM MgCl_2_, pH 8). The beads were resuspended in 100 µL benzonase buffer supplemented with protease inhibitor and 25 U of Benzonase nuclease (Sigma). The samples were incubated at 37 °C for 10 min, while shaking at 900 rpm. After this, the beads were washed once with high-salt NT-P buffer and twice with FastAP buffer (10 mM Tris-HCl, 5 mM MgCl_2_, 100 mM KCl, pH 8). The beads were resuspended in 150 µL of FastAP buffer supplemented with protease inhibitor and 2 U of Fast Alkaline Phosphatase (Thermo). Samples were incubated at 37 °C for 30 min, shaking at 900 rpm. The beads were washed once with high-salt NT-P buffer and twice with PNK buffer (50 mM Tris-HCl, 10 mM MgCl_2_, 0.1 mM spermidine, pH 8). The magnetic beads were resuspended in 100 µL of PNK buffer supplemented with protease inhibitor, 10 µCi of [γ-^32^P]-ATP and 1 µL of PNK (Thermo). Samples were incubated at 37 °C for 30 min, while shaking at 900 rpm. The beads were washed once with high-salt NT-P buffer and twice with NT-P buffer. For the elution of the protein-RNA complexes, the beads were resuspended in 35 µL of TBS buffer (50 mM Tris-HCl, 150 mM NaCl, pH 7.5) supplemented with 0.5 µg/µL FLAG peptide (Sigma) and incubated at 4 °C for 15 min, while gently shaking at 900 rpm. After separation of beads to the supernatant using a magnetic rack, protein loading dye was added to supernatant (5x concentrated loading dye: 25 mM Tris-HCl pH 6.8, 25% glycerol, 10% SDS, 50 mM DTT and 0.05% bromophenol blue). After 5 min of denaturation at 95 °C, the samples were separated on 15% SDS-PAA gels and electroblotted onto nitrocellulose membranes (Amersham). Radioactive ink was added to the corners of the membrane to facilitate the overlay of the size markers. Radioactive signals were detected using a Typhoon FLA 9500 (GE Healthcare). After detection of the radioactive signal, the membranes were blocked in 5% skim milk in T-PBS for 1 h and subsequently used for Western blotting using anti-FLAG antibodies as described above.

### Growth assay and time-course expression of 3xFLAG-PatR

Cells from liquid cultures of *Nostoc*, Δ*patR* and Δ*patR*+ were grown in BG11 medium under standard conditions as explained above. After 5 d, they were collected and resuspended in BG11_0_ at OD_750_ = 0.1. Five-fold serial dilutions were prepared and 10 μL of each dilution was plated on BG11_0_ plates containing ammonium (2.5 mM NH_4_ and 12 mM N-tris (hydroxymethyl) methyl-2-aminoethanesulfonic acid-NaOH buffer (pH 7.5), nitrate (17 mM NO_3_) or lacking combined nitrogen (N_2_). Growth was analyzed after 13 d of incubation at 30 °C. To analyze heterocyst differentiation by optical microscopy, liquid cultures of *Nostoc*, Δ*patR* and Δ*patR*+ growing photoautotrophically in BG11 under standard conditions were harvested, washed and resuspended in media without combined nitrogen source (BG11_0_) as described above. After 24 of cultivation, samples of Δ*patR* were taken and heterocysts were stained with alcian blue as previously described^85^. After 7 d of cultivation, visualization of heterocysts was not possible in Δ*patR* because this strain did not grow diazotrophically.

Liquid cultures of the strain 3xFLAG-PatR were grown in BG11 under standard conditions, cells were harvested by centrifugation, and heterocyst differentiation was induced as explained above. Samples were taken at different time points after combined nitrogen removal. Cells were mechanically lysed as explained above and 30 µg of total soluble protein was separated on a 15% PAA-SDS gel and transferred to a nitrocellulose membrane. The membrane was stained with Ponceau Red (0.1% (w/v) in 5% acetic acid) as loading control, blocked in 5% skim milk in T-PBS for 1 h and subsequently used for Western blotting using anti-FLAG antisera as described above.

### Microscopy and image analysis

Filaments stained with alcian blue were visualized using an Olympus BX60 microscope. Fluorescence of *Nostoc* filaments carrying plasmid pMBA118 growing for 5 d on top of solidified BG11 or BG11_0_ (nitrogen-free) medium supplemented with appropriate antibiotics was analyzed by a Nikon CFI Plan apochromat VC 60x 1.4 oil immersion objective and DIC N2 condenser attached to a Nikon Eclipse T1 confocal microscope. The samples were excited at 488 nm and the fluorescent emission was filtered by a HQ filter set (GFP, 500-550 nm window) and a Cy-5 filter set (photosynthetic pigment autofluorescence, 663-738 nm window). For quantification purposes, all images were taken with the same settings. Fluorescence of WT and *ΔpatR* filaments carrying plasmids pSAM301 or pSAM270 growing for 5 d on top of solidified BG11 or BG11_0_ (nitrogen-free) medium were analyzed using a Leica HCX PLAN-APO 63× 1.4 NA oil immersion objective attached to a Leica TCS SP2 laser-scanning confocal microscope. Samples were excited at 488 nm by an argon-ion laser and the fluorescent emission was monitored by collection across windows of 500-538 nm (GFP) and 630-700 nm (photosynthetic pigment autofluorescence). For quantification purposes, the WT and *ΔpatR* strains were grown in the sample plate and images were taken with the same settings. Images were analyzed with Fiji^86^.

### Phylogenetic analysis of PatR and conservation of *patR* promoter sequences

A BLASTP search was performed using default parameters and the *Nostoc* PatR protein sequence (WP044521042.1) as a query. Because 1,658 homologs were found, we decided to use only 107 homologs from genomes associated with reference cyanobacteria^87^ for the downstream analysis. Multiple alignment was performed using ClustalW. The best distance model was selected using MEGAX^88^. The evolutionary history was inferred by using the Maximum Likelihood method and Jones et al. w/freq. model^89^. 1,000 iterations of bootstrapping were carried out to infer the confidence of the branches. Initial tree(s) for the heuristic search were obtained automatically by applying Neighbor-Join and BioNJ algorithms to a matrix of pairwise distances estimated using the JTT model, and then selecting the topology with the higher log likelihood value. A discrete Gamma distribution was used to model evolutionary rate differences between sites (5 categories; +G, parameter = 1.1415). The rate variation model allowed for some sites to be evolutionarily invariable ([+I], 1.40% sites).

### Structural analysis

Alphafold prediction for PatR (AF-Q8YWB5-F1), Slr1122 (AF-P72645-F1) and PMM0486 (AF-Q7V2J0-F1) were downloaded from the AlphaFold Protein Structure Database. Sequences for the conserved domains of PatR (residues 70-158 and residues 198-279, respectively) were folded by the ESMFold method and compared with the structure of other proteins through the Foldseek webserver^44^. The visualization of the PatR structure, the calculation of the Coulombic electrostatic potential (ESP) and the superimposition of structural homologs were performed by ChimeraX^90^.

## Data availability

Source data are provided as a Source Data file. MS raw data is deposited at the ProteomeXchange Consortium (http://proteomecentral.proteomexchange.org) via the PRIDE partner repository^91^ under the identifier: PXD050404. All code involving the bioinformatics workflow and the shiny app is available on Github (https://github.com/manbrealv/R-DeeP-Nostoc). A shiny app was programmed to facilitate the accessibility and visualization of the data: https://sunshine.biologie.uni-freiburg.de/R-DeeP-Nostoc/.

## Acknowledgments

The authors thank Prof. Enrique Flores and Prof. Antonia Herrero for sharing the *ΔhetR* CSSC2 strain. We also thank Marc Broghammer for help in cloning the Asl3888-3xFLAG fusion and Karsten Voigt and Edith Ams for help in the implementation of the shiny app on the webserver.

## Author contributions

MBA designed the project. MBA, AMP, MMS and WRH secured funding. The R-DeeP approach, PNK assays, phylogenetic analysis and construction of the mutants and reporter strains was performed by MBA. DP, CP, FS and MMS performed and interpreted the proteomic analysis. Bioinformatic analysis of R-DeeP data was performed by FS, DP and MBA. The modified version of TriPepSVM was implemented by HRR and MBA. Interpretation of the R-DeeP data and SVM output was performed by MBA and WRH. Growth analysis and microscopy experiments for *patR* were performed and interpreted by AV, AMP and MBA. MBA and WRH wrote the manuscript with the input of all authors.

