## Supplementary_Figures_and_Tables for "RNA-binding proteins identified by R-DeeP/TripepSVM are involved in heterocyst differentiation"

**Content:**

**Supplementary Figures:** p.2
**Supplementary Tables:** p.10
**Overview on Supplemental Datasets:** p.17
**Supplemental references:** p.18

### Fig. S1

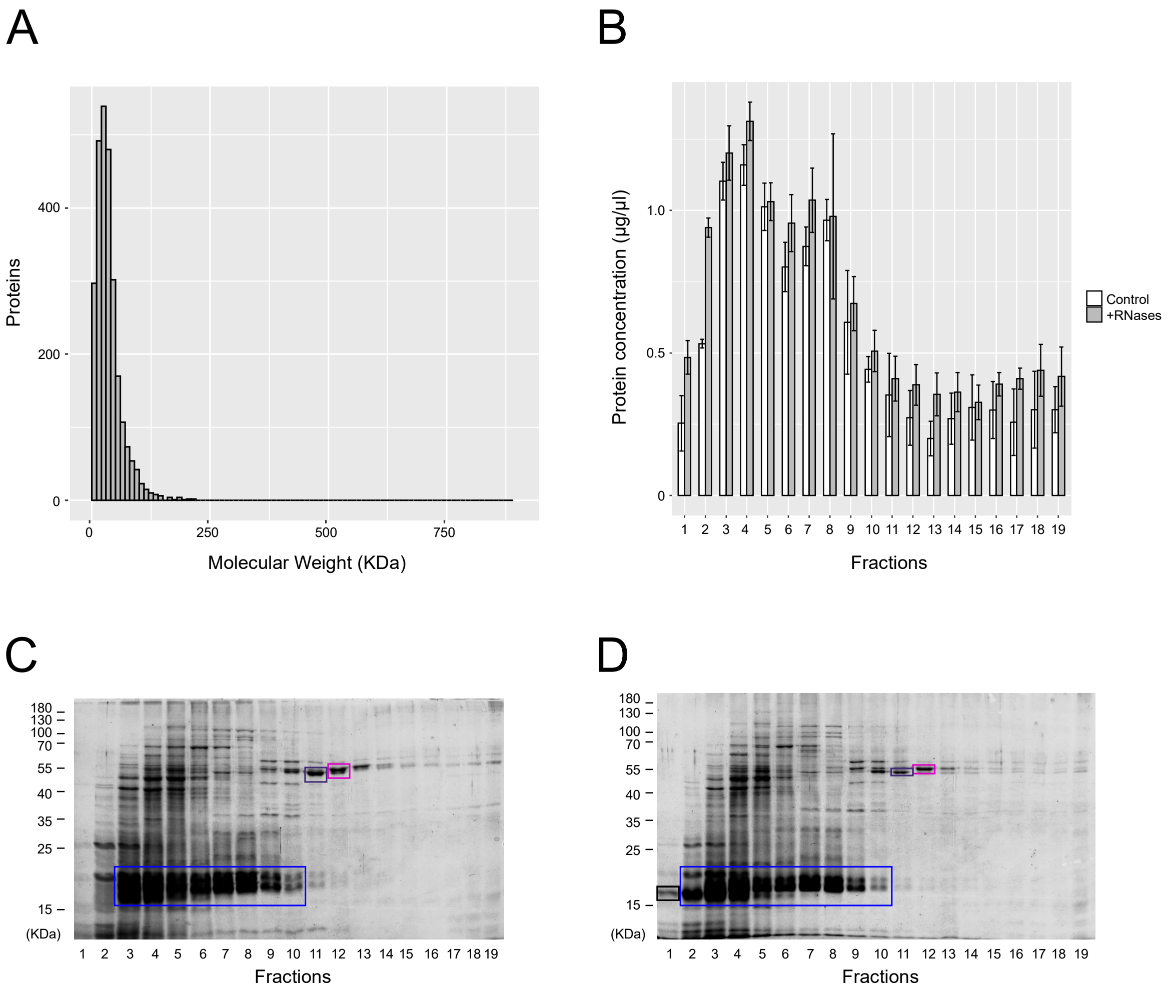


**Figure S1 | Global overview of protein distribution in the density gradients. a** Histogram of the theoretical distribution of the identified proteins in monomeric form. A limit of the x-axis from 0 to 900 KDa is plotted because this is the resolution of a 10-40% sucrose gradient. **b** Histogram of protein concentration in the 19 fractions. The data are presented as the mean ± standard deviation of the three replicates of each fraction under the different conditions. **c-d** Coomasie-stained 15% polyacrilamide SDS-PAGE gels with fractions from a representative gradient out of three from a control sample (c) or RNAse-treated sample (d). 35 µl of each fraction was loaded. Some highly expressed proteins are highlighted; blue frame (proteins from phycobilisomes), purple frame (large subunit of RuBisCO, RbcL), pink frame (glutamine synthetase, GlnA) and black frame (RNAses added to the treated samples). The position of size markers (in kilodaltons) is indicated on the left side.

### Fig. S2

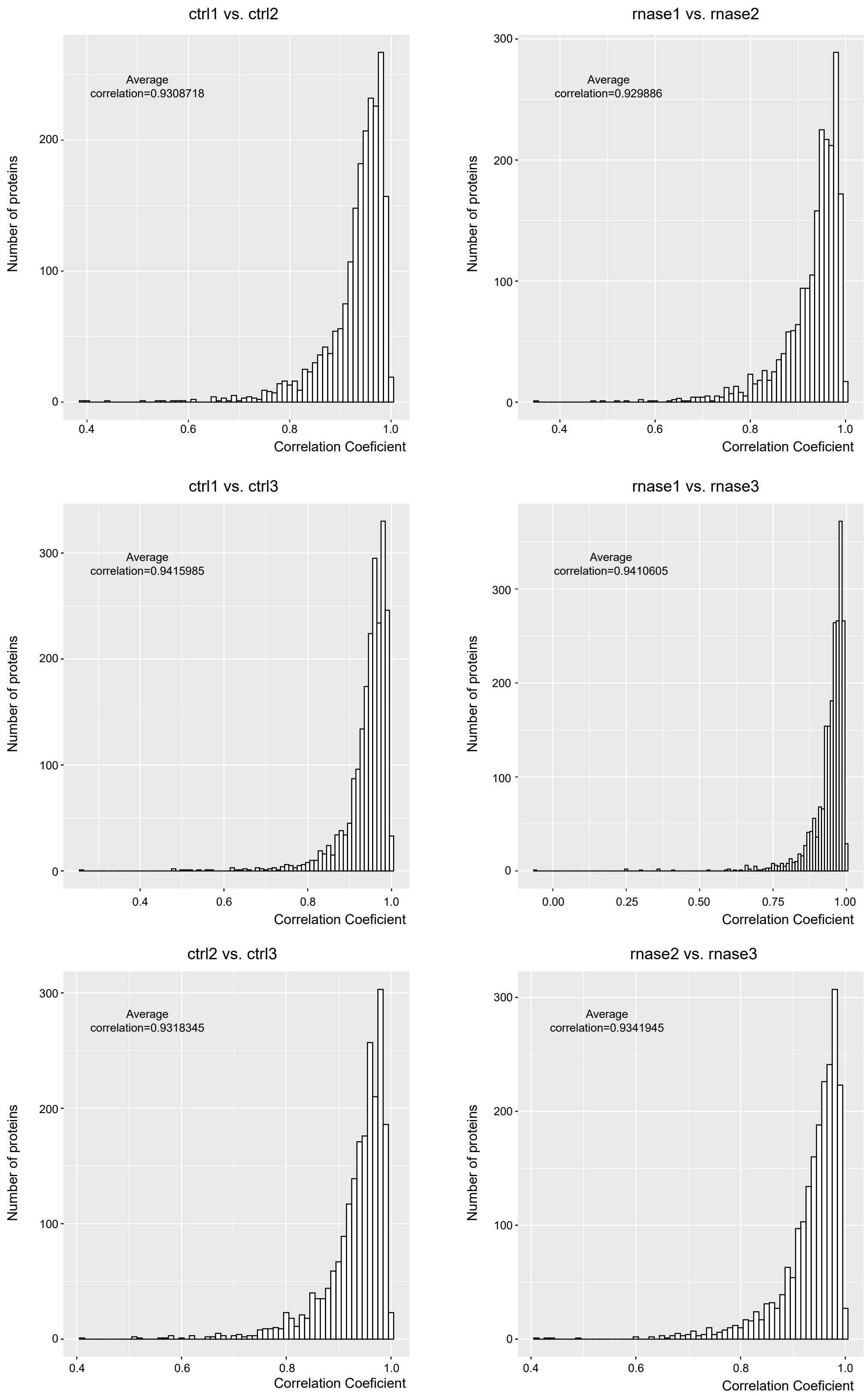


**Figure S2 | Correlation between samples.** Histograms of Spearman correlation coefficients from the comparison between control replicates or RNase-treated replicates.

### Fig. S3

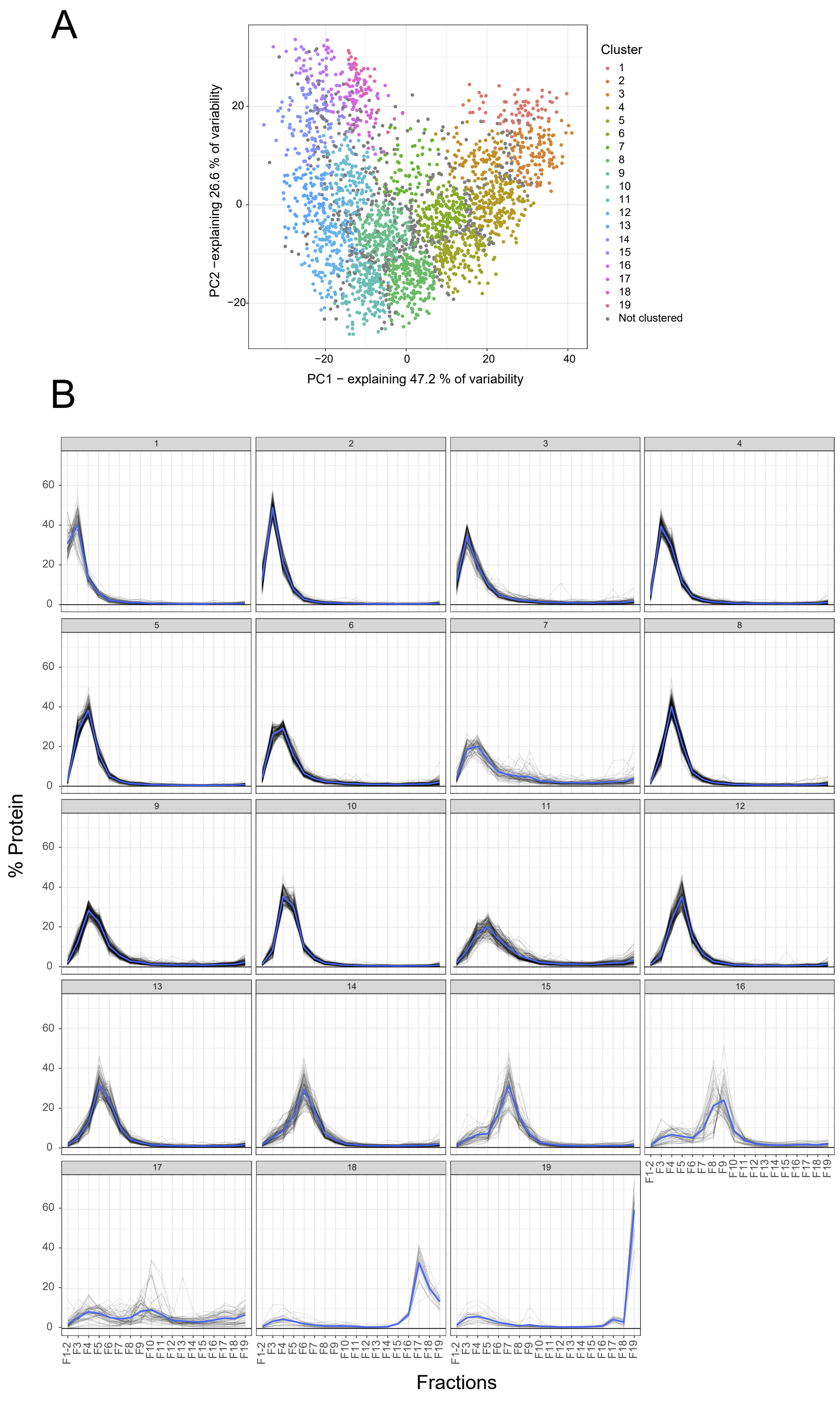


**Figure S3 | Data clustering of control gradients. a** PCA plot of the mean protein profiles between the three control gradients. The variability explained by the first two principal components is shown. The results of the soft clustering analysis are shown in colour. Because only proteins with a confidence greater than 90% are considered to be clustered, some proteins that do not clearly belong to any cluster are kept as “not clustered”. **b** Line plots of the average distribution of proteins in the 19 clusters. Grey lines show the distribution of individual proteins. The blue line shows the mean distribution of each cluster.

### Fig. S4

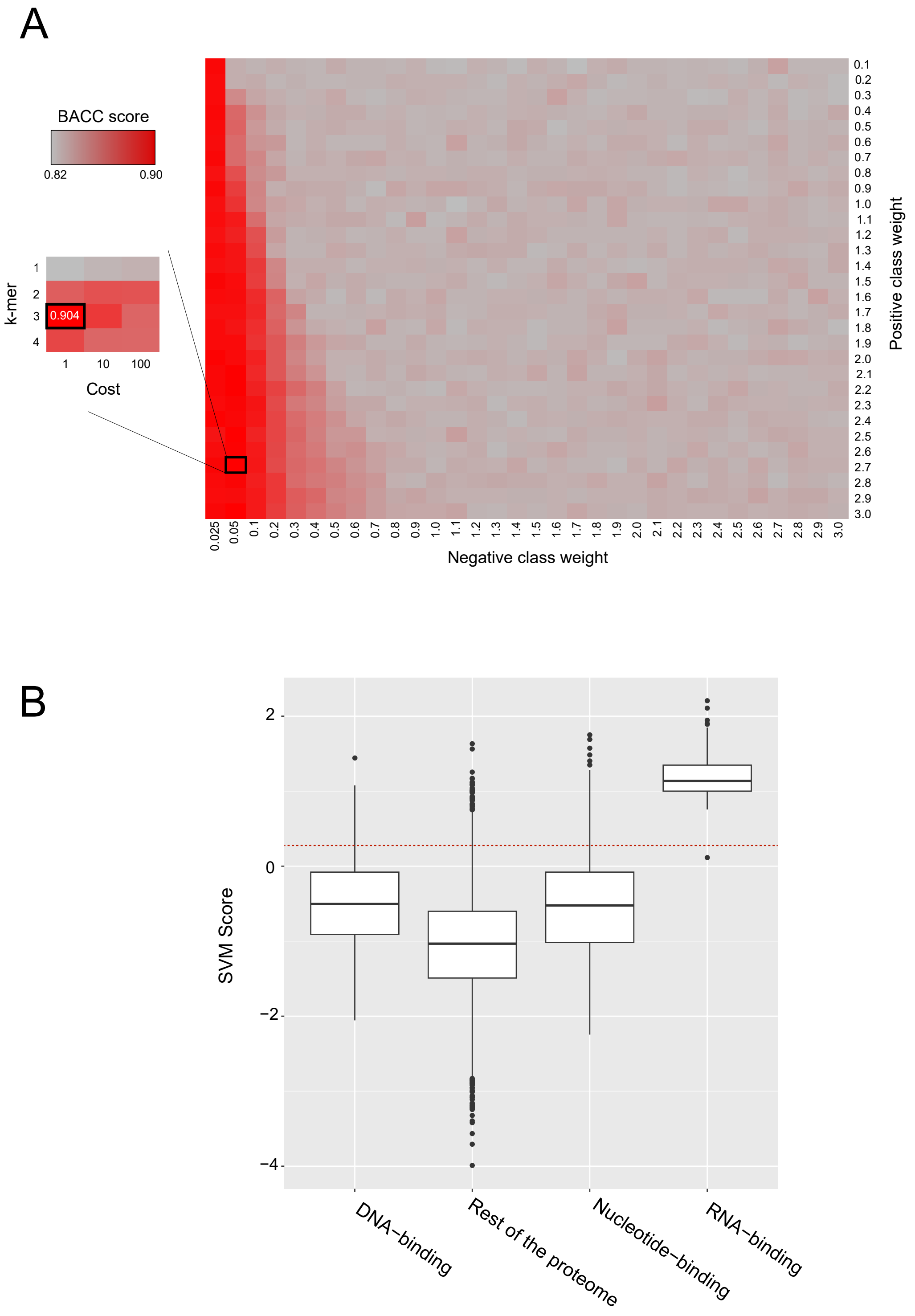


**Figure S4 | Parameter optimisation of the modified TriPepSVM approach. a** Heatmap showing the balanced accuracy score (BACC) for a grid search using a 10-k-fold cross-validation for a combination of parameters: positive class weight, negative class weight, k-mers and cost. The BACC score for the best combination of parameters is highlighted with a black frame. b Threshold selection for predicting RBPs using the SVM score. The box plot shows the distribution of the SVM score for previously known DNA-binding, RNA-binding and nucleotide-binding proteins according to the QuickGO annotation. The selected threshold of 0.25 is indicated by a red dashed line.

### Fig. S5

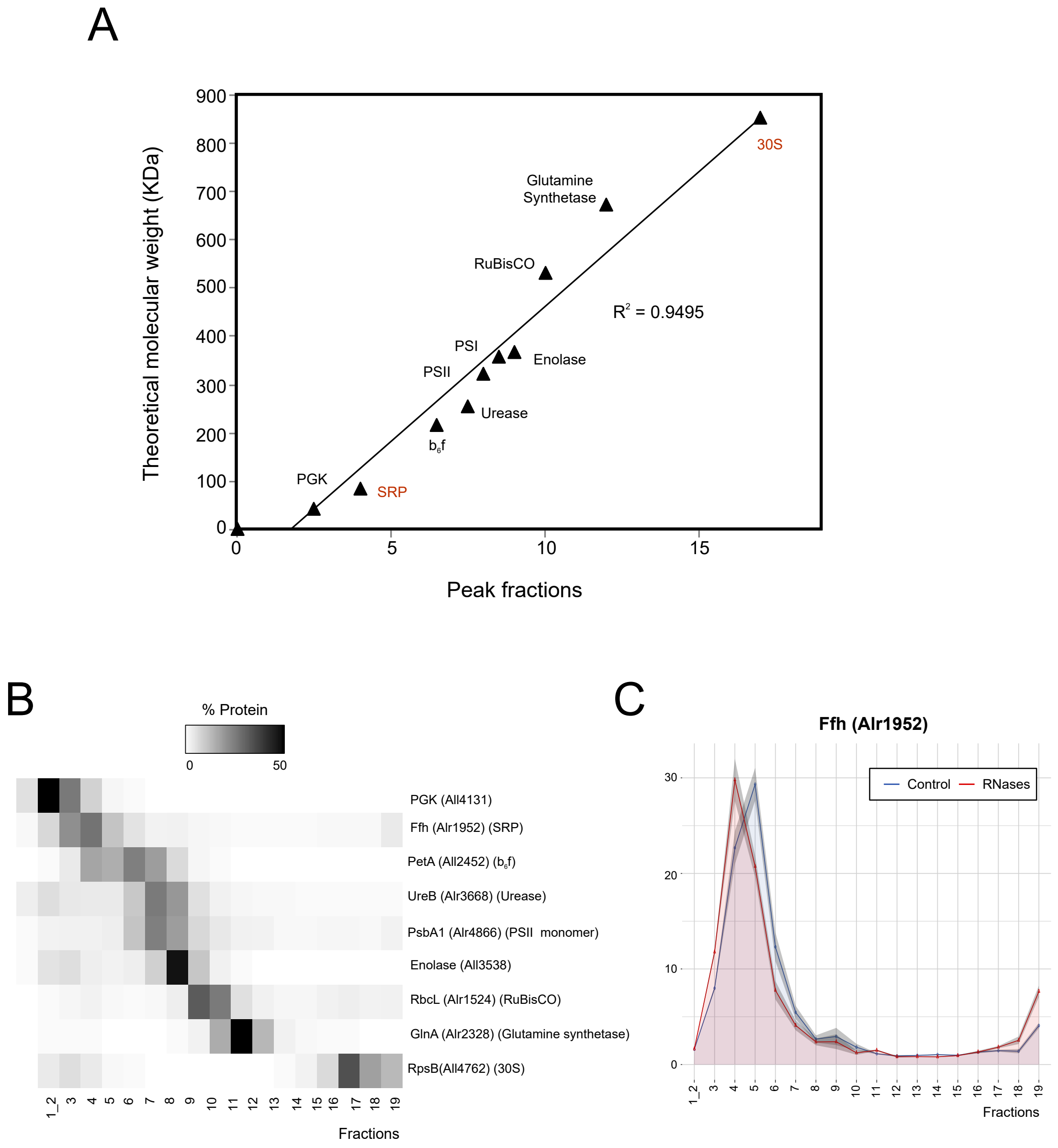


**Figure S5 | Sedimentation velocity in comparison to molecular weight and resolution limit of gradients. a** Selected protein complexes (black triangles) illustrate the correlation between molecular weight and sedimentation velocity of proteins in the untreated gradients. Protein-protein complexes are shown in black, while RNA-protein complexes are shown in red. The peak fraction of a representative protein of each complex was selected for the calibration curve. **b** Heatmap of the mean normalised abundance of the representative proteins used for the calibration curve of a. The sum of a protein abundance along the gradient is normalised to 100%. **c** Limitations in the resolution of the gradients.The protein subunit of the signal recognition particle (Ffh) is shown as an example of an RBP that interacts with a small RNA. The weight loss for the SRP after 4.5S RNA degradation is so small that, although there is a shift, it is not detected as statistically significant by our approach. The graph shows the normalised protein abundance measured by MS along the 18 analysed fractions. The sum of protein abundance along the gradient is normalised to 100%. Raw data (mean of 3 replicates) are indicated by the lines. The shading indicates the standard deviation between replicates.

### Fig. S6

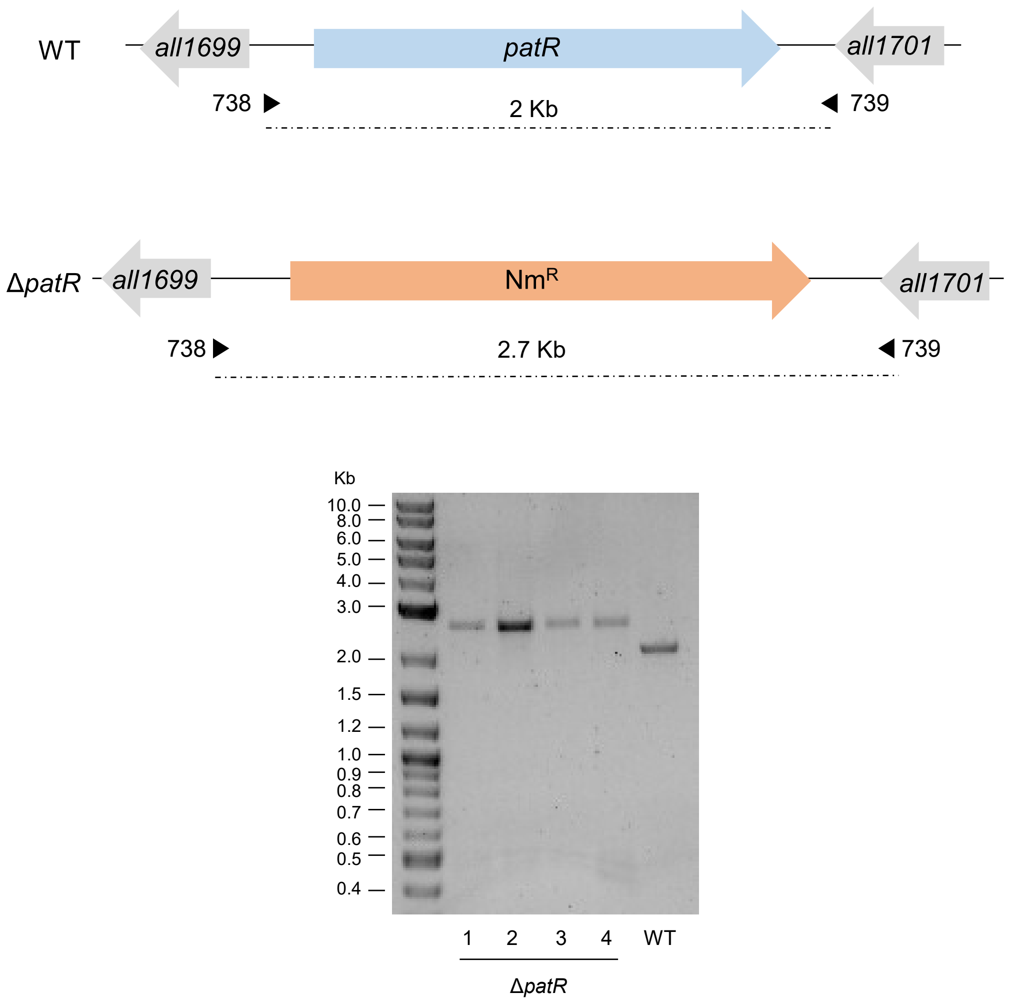


**Figure S6 | Genomic analysis of Δ*patR*.** A scheme of the regions surrounding *patR* is shown for the WT and the Δ*patR* mutant. DNA fragments were amplified by PCR using as template DNA from 4 clones of Δ*patR* or WT and oligonucleotides 738 and 739.

### Fig. S7

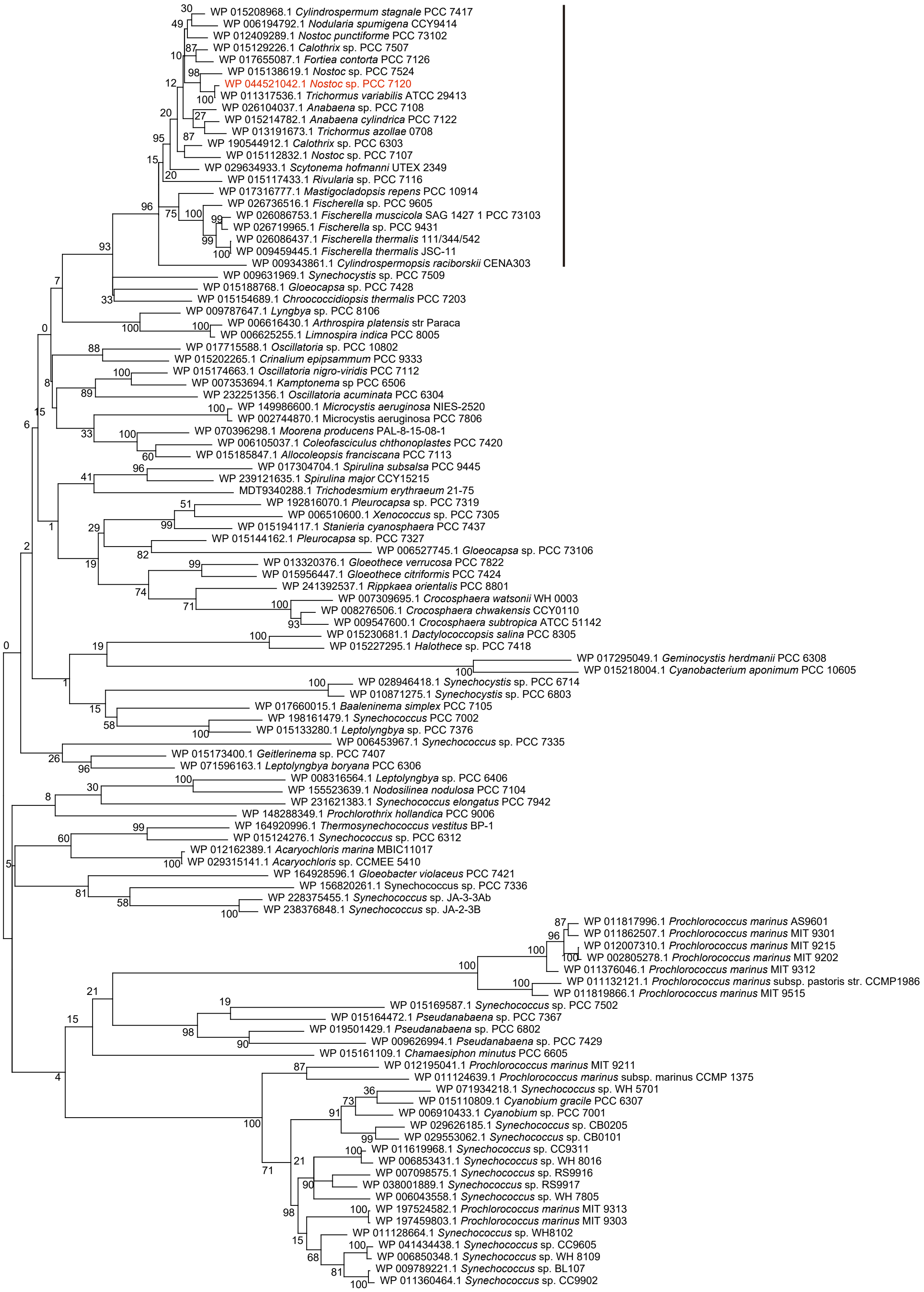


**Figure S7 | Phylogenetic tree of PatR.** The evolutionary history was inferred by using the Maximum Likelihood method and Jones et al. w/freq. model[1]. The tree with the highest log likelihood (-40697.75) is shown. 1000 iterations of bootstraping were carried out to infer the confidence of the branches. The percentage of trees in which the associated taxa clustered together is shown next to the branches. Initial tree(s) for the heuristic search were obtained automatically by applying Neighbor-Join and BioNJ algorithms to a matrix of pairwise distances estimated using the JTT model, and then selecting the topology with superior log likelihood value. A discrete Gamma distribution was used to model evolutionary rate differences among sites (5 categories (+G, parameter = 1.1415)). The rate variation model allowed for some sites to be evolutionarily invariable ([+I], 1.40% sites). The tree is drawn to scale, with branch lengths measured in the number of substitutions per site. Evolutionary analyses were conducted in MEGA X[2]. Vertical black line shows the heterocyst-forming cyanobacteria.

### Fig. S8

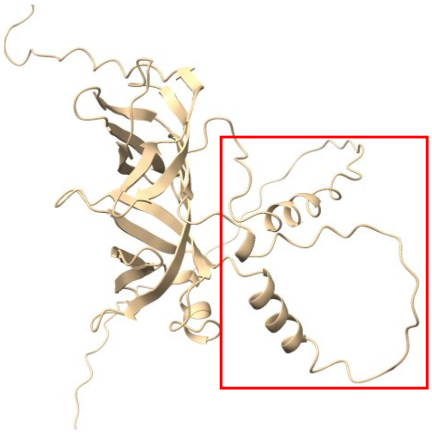

PatR
(*Nostoc* sp.PCC 7120)

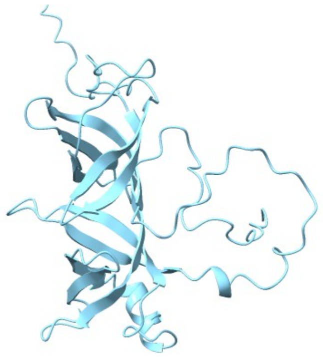

Slr1122
(*Synechocystis* sp. *PCC 6803*)

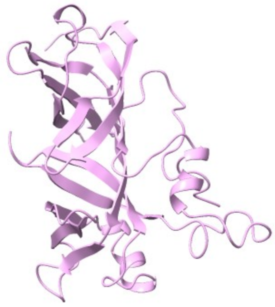

PMM0486
(*Prochlorococcus marinus* MED4)

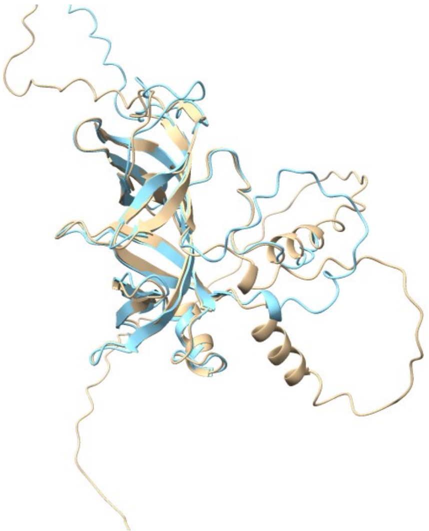

PatR-Slr1122

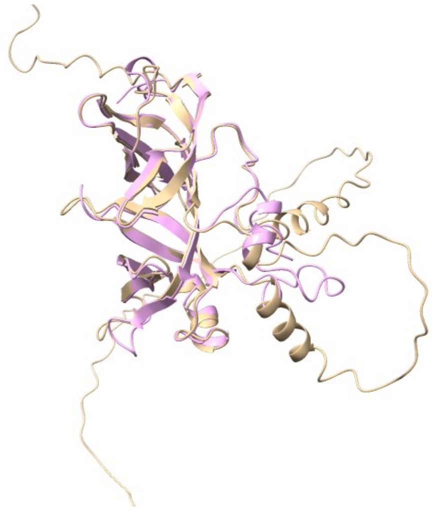

PatR-PMM0486

**Figure S8 | Structures of PatR, Slr1122 and PMM0486.** Structures were downloaded from AlphaFold Protein Structure Database; PatR (AF-Q8YWB5-F1), Slr1122 (AF-P72645-F1) and PMM0486 (AF-Q7V2J0-F1). The superimposition of the structural homologs was performed using the matchmaker function of ChimeraX. The two helixes and loop conserved in heterocyst-forming cyanobacteria are framed in red.

**Table S1.** Proteomes selected for the assembly of positive and negative training datasets for the SVM approach.

| Organism | Taxonomy ID UniProt | RBP | NRBP |
|---|---|---|---|
| *Nostoc* sp. PCC 7120 | 103690 | 107 | 4741 |
| *Synechocystis* sp. PCC 6803 | 1111708 | 112 | 2568 |
| *Nostoc punctiforme* PCC 73102 | 63737 | 108 | 5082 |
| *Trichodesmium erythraeum* IMS101 | 203124 | 99 | 3144 |
| *Cyanothece* sp. PCC 7425 | 395961 | 108 | 3969 |
| *Nodularia spumigena* CCY9414 | 313624 | 107 | 3524 |
| *Nostoc azollae* 0708 | 551115 | 99 | 2562 |
| *Fischerella* sp. NIES-3754 | 1752063 | 108 | 3727 |
| *Rivularia* sp. PCC 7116 | 373994 | 111 | 5077 |
| *Prochlorococcus marinus* subsp. pastoris CCMP1986 | 59919 | 86 | 1337 |
| *Escherichia coli* K-12 | 83333 | 182 | 3201 |
| *Salmonella typhimurium* LT2 | 99287 | 139 | 3793 |

For each proteome is shown the number of RBPs annotated in QuickGO (RBP) used as positive training dataset and the number of proteins treated as negative dataset (NRBP). The Taxonomy ID of each genome is also included.

**Table S2.** Genomes used for the homolog search for *Nostoc* proteome.

| Genome | IMG Genome ID |
|---|---|
| *Trichodesmium erythraeum* IMS101 | 2639763089 |
| *Arthrospira platensis* NIES-39 | 650377906 |
| *Oscillatoria acuminata* PCC 6304 | 2509276028 |
| *Oscillatoria nigro-viridis* PCC 7112 | 2503982035 |
| *Fischerella* sp. NIES-3754 | 2687453106 |
| *Fischerella* sp. NIES-4106 | 2775506903 |
| *Calothrix parietina* PCC 6303 | 2503982036 |
| *Nostoc* sp. PCC 7524 | 2509601032 |
| *Nostoc* sp. PCC 7120 | 2914518388 |
| *Trichormus variabilis* ATCC 29413 | 646564504 |
| *Nostoc* sp. PCC 7107 | 2503707008 |
| *Cylindrospermum stagnale* PCC 7417 | 2509601025 |
| *Anabaena cylindrica* PCC 7122 | 2503982047 |
| *Cylindrospermopsis raciborskii* Cr2010 | 8001103026 |
| *Trichormus azollae* 0708 | 648028001 |
| *Nodularia spumigena* CCY9414 | 639857037 |
| *Microchaete diplosiphon* NIES-3275 | 2775506820 |
| *Calothrix* sp. PCC 7507 | 2505679032 |
| *Nostoc punctiforme* PCC 73102 | 2617270889 |
| *Tolypothrix* sp. PCC 7910 | 2883284429 |
| *Rivularia* sp. PCC 7116 | 2510065008 |
| *Gloeocapsa* sp. PCC 7428 | 2503754017 |
| *Chroococcidiopsis thermalis* PCC 7203 | 2503538021 |
| *Dactylococcopsis salina* PCC 8305 | 2509276056 |
| *Spirulina major* PCC 6313 | 2506520014 |
| *Cyanothece* sp. PCC 7425 | 643348534 |
| *Synechocystis* sp. PCC 6803 | 2514885032 |
| *Pleurocapsa minor* PCC 7327 | 2509276061 |
| *Microcystis aeruginosa* NIES-843 | 641522640 |
| *Microcystis aeruginosa* PCC 7806SL | 2751185885 |
| *Leptolyngbya* sp. PCC 7376 | 2503754048 |
| *Cyanobacterium stanieri* PCC 7202 | 2503283023 |
| *Cyanobacterium aponinum* PCC 10605 | 2503707009 |
| *Stanieria cyanosphaera* PCC 7437 | 2503754019 |
| *Microcoleus* sp. PCC 7113 | 2509276031 |
| *Crinalium epipsammum* PCC 9333 | 2504643013 |
| *Chamaesiphon minutus* PCC 6605 | 2510436000 |
| *Geitlerinema* sp. PCC 7407 | 2503538020 |
| *Prochlorococcus marinus* MIT9211 | 641228501 |
| *Prochlorococcus marinus* MIT9303 | 640069323 |
| *Prochlorococcus marinus* MIT9313 | 637000211 |
| *Prochlorococcus marinus* NATL1A | 640069325 |
| *Prochlorococcus marinus pastoris* CCMP 1986 | 637000214 |
| *Synechococcus* sp. CC9311 | 637000309 |
| *Synechococcus* sp. WH 8103 | 2687453380 |
| *Synechococcus* sp. WH 8109 | 2563366603 |
| *Cyanobium gracile* PCC 6307 | 2508501011 |
| *Synechococcus elongatus* PCC 6301 | 637000307 |
| *Synechococcus elongatus* PCC 7942 | 637000308 |
| *Thermosynechococcus elongatus* BP-1 | 637000320 |
| *Acaryochloris marina* MBIC11017 | 641228474 |
| *Pseudanabaena* sp. PCC 7367 | 2504643012 |
| *Synechococcus* sp. JA-2-3B'a (2-13) | 637000312 |
| *Synechococcus* sp. JA-3-3Ab | 637000313 |
| *Gloeobacter violaceus* PCC 7421 | 637000121 |

**Table S3.** Strains.

| Strain | Description |
|---|---|
| ***Escherichia coli*** | |
| DH5α | Used for routine transformation[3] |
| ***Nostoc* sp.** | |
| PCC 7120 | Wild type (Pasteur Culture Collection) |
| CSSC2 | $Nm^R$, Δ*hetR* mutant[4] |
| HetR-3xFLAG | $Nm^s$, replacement of the Nm resistance cassette of CSSC2 Δ*hetR* mutant with a native copy of *hetR* fused to 3xFLAG. Complementation carried out through pMBA160 insertion (this work). |
| Δ*patR* | $Nm^R$, Δ*patR* mutant. Replacement of *patR* locus by a CK3 cassette (neomycin resistance cassette) carried out by plasmid pMBA66 (this work). |
| Δ*patR* + | $Nm^RSm^RSp^R$. Insertion of pMBA79 in alpha plasmid of Δ*patR*. Complementation of the knock-out mutant with a copy of *patR* expressed from *rnpB* promoter (this work). |
| RnpA-3xFLAG | $Sm^RSp^R$. Insertion of pMBA189 in *rnpA* locus of WT. Expression of RnpA fused to 3xFLAG under native promoter (this work). |
| Alr2890-3xFLAG | $Sm^RSp^R$. Insertion of pMBA213 in *alr2890* locus of WT. Expression of Alr2890 fused to 3xFLAG under native promoter (this work). |
| PrpA-3xFLAG | $Sm^RSp^R$. Insertion of pMBA180 in *prpA(alr3731)* locus of WT. Expression of PrpA fused to 3xFLAG under native promoter (this work). |
| Asl3888-3xFLAG | $Sm^RSp^R$. Insertion of pMB2 in alpha plasmid of WT. Expression of Asl3888 fused to 3xFLAG under native promoter (this work). |
| VIPP-3xFLAG | $Sm^RSp^R$. Insertion of pMBA171 in *vipp1(all2342)* locus of WT. Expression of Vipp1 fused to 3xFLAG under native promoter (this work). |
| TrpB-3xFLAG | $Sm^RSp^R$. Insertion of pMBA183 in *trpB (all0410)* locus of WT. Expression of TrpB fused to 3xFLAG under native promoter (this work). |
| Alr2809-3xFLAG | $Sm^RSp^R$. Insertion of pMBA167 in *alr2809* locus of WT. Expression of Alr2809 fused to 3xFLAG under native promoter (this work). |
| PatU3-3xFLAG | $Sm^RSp^R$. Insertion of pMBA178 in *patU3 (alr0101)* locus of WT. Expression of PatU3 fused to 3xFLAG under native promoter (this work). |
| 3xFLAG-PatR | $Nm^RSm^RSp^R$. Insertion of pMBA149 in alpha plasmid of Δ*patR*. Complementation of the knock-out mutant with a fusion of 3xFLAG to a native copy of *patR* under the native promoter of *patR*. (This work). |
| $P_{patR}$:GFP | $Sm^RSp^R$, pMBA118 inserted in plasmid alpha. *gfpmut2* under the control of the *patR* promoter (this work). |
| $P_{nsiR1}$:GFP/ Δ*patR* | $Nm^RSm^RSp^R$. Insertion of pSAM301 in alpha plasmid of Δ*patR*. *gfpmut2* under the control of the *nsiR1* promoter (this work). |
| $P_{hetR}$:GFP/ Δ*patR* | $Nm^RSm^RSp^R$. Insertion of pSAM270 in alpha plasmid of Δ*patR*. *gfpmut2* under the control of the *hetR* promoter (this work). |
| $P_{nsiR1}$:GFP/ WT | $Nm^RSm^RSp^R$. Insertion of pSAM301 in alpha plasmid of WT. *gfpmut2* under the control of the *nsiR1* promoter[5]. |
| $P_{hetR}$:GFP/ WT | $Nm^RSm^RSp^R$. Insertion of pSAM270 in alpha plasmid of WT. *gfpmut2* under the control of the *hetR* promoter[6]. |

**Table S4.** Oligonucleotides.

| ID | Sequence | Description |
|---|---|---|
| 90 | **TAATACGACTCACTATAGGG**CTCTCTGATAGCGGAACTGG | DNA template for *rnpB* RNA probe |
| 91 | CCGCCAAGATTGGGGACTGGGG | |
| 75 | TATAGCGGAGACGCATGTTTCCGTT | DNA template for *yfr1* RNA probe |
| 76 | **TAATACGACTCACTATAGGG**ATGTAATACATAAGGAACCGCCC | |
| 77 | **TAATACGACTCACTATAGGG**TTTTACCCCGTCGAAGTAGCT | DNA template for *nsiR8* RNA probe |
| 78 | GGGAATTGTGAGGATAGAAATGATT | |
| 20 | AGCCAGCTAGGGGAGTTAGTTATCAGGGTGCATCTACC | Oligonucleotide *nsiR1.4* probe |
| 111 | CCGGTCCTGACAAGGTTTAGGCGTTGC | Oligonucleotide 4.5S probe |
| 5S | TAGCAGCGTTTCACCTCTGAGTTCGG | Oligonucleotide 5S probe |
| 669 | GTTTTGGATCCCTTGGTTACTAGCACTATTC | Δ*patR* generation |
| 670 | CCTTCCTCGAGATCCCAATTATGGGATTTTA | |
| 671 | GGGATCTCGAGGAAGGGGGCAGGGAAGTAGG | |
| 672 | GTTTTGGATCCCGTTGGGTGAGGGGTTGTTA | |
| 738 | GTTTTATCGATGATTAGAGGCGAGCGAGTCGC | Expression of *patR* under *rnpB* promoter. |
| 739 | GTTTTCTCGAGGATGCCAAGCAAATGGATGCT | |
| 32 | GTTTGAGCTCTTATTTGTAGAGTTCATCCATGCC | 3xFLAG-sfGFP fusion in pMBA132 |
| 34 | GTTTATGCATGATTATAAAGATCATGATGGCGATTATAAAGATC ATGATATTGATTATAAAGATGATGATGATAAACTCGAGGGATCC GCTGGCTCCGCTGCT | |
| 33 | GTTTATGCATATGAGCAAAGGAGAAGAACTT | sfGFP-3xFLAG fusion in pMBA132 |
| 35 | GTTTGAGCTCTTATTTATCATCATCATCTTTATAATCAATATCAT GATCTTTATAATCGCCATCATGATCTTTATAATCCTCGAGTTTGT AGAGTTCATCCATGCCATG | |
| 129 | GTTTTCTGCAGCTTCTGTACAAGTCAAACCAGAAAG | Alr2809-3xFLAG fusion in pMBA132 |
| 130 | GTTTTCTCGAGCCCCATCTGTCCTCGTACCCATTC | |
| 136 | GTTTTCTGCAGTCACTCAAATGGGTAGCGACGA | VIPP1-3xFLAG fusion in pMBA132 |
| 137 | GTTTTCTCGAGCAACTGATCTAATTGTTTGCGTAGG | |
| 148 | GTTTTCTGCAGATTCCTACTGTGCAAGAACG | PatU3-3xFLAG fusion in pMBA132 |
| 149 | GTTTTCTCGAGTGTTGCGGGATTAATGACAAAT | |
| 151 | GTTTCTGCAGTTGAGGATGTATTAGGTAAGCGC | PrpA-3xFLAG fusion in pMBA132 |
| 152 | GTTTTCTCGAGCAGTCGTGGATCTACGGGTTC | |
| 156 | GTTTCTGCAGGTGGTAAGCACTCCAGAAATTA | TrpB-3xFLAG fusion in pMBA132 |
| 157 | GTTTTCTCGAGATAATTAAGGACTTTGGCGACG | |
| 166 | GTTTTCTGCAGTTCCTTTTTGATTGAAACTTGAGC | RnpA-3xFLAG fusion in pMBA132 |
| 167 | GTTTTCTCGAGCGAATGCCCATGTAATACCT | |
| 194 | GTTTTCTGCAGCGCTGATACAAGTCCTGTGTG | Alr2890-3xFLAG fusion in pMBA132 |
| 195 | GTTTTCTCGAGCATATACCTGGTGAGTTGGACGC | |
| P03 | GTTTCTCGAGTTCAGCATCAACTGCTGCTGCG | Asl3888-3xFLAG fusion in pMBA132 |
| P04 | GTTTCTGCAGGTTTCAATCCCCGACTCCAACTG | |
| 131 | GTTTTGAGCTCCTTCTGTACAAGTCAAACCAGAAAG | Alr2809-3xFLAG fusion in pCSV3 |
| 138 | GTTTTGAGCTCTCACTCAAATGGGTAGCGACGA | VIPP1-3xFLAG fusion in pCSV3 |
| 150 | GTTTTGAGCTCATTCCTACTGTGCAAGAACG | PatU3-3xFLAG fusion in pCSV3 |
| 153 | GTTTTGAGCTCTTGAGGATGTATTAGGTAAGCGC | PrpA-3xFLAG fusion in pCSV3 |
| 158 | GTTTTGAGCTCGTGGTAAGCACTCCAGAAATTA | TrpB-3xFLAG fusion in pCSV3 |
| 168 | GTTTTGAGCTCTTCCTTTTTGATTGAAACTTGAGC | RnpA-3xFLAG fusion in pCSV3 |
| 196 | GTTTTGAGCTCCGCTGATACAAGTCCTGTGTG | Alr2890-3xFLAG fusion in pCSV3 |
| 19 | AGCTGTCAATGTCTACCACTTT | Cloning of protein fusions in pCSV3 |
| 57 | GTTTTCTGCAGCCAATAATGGGAACAACTGAAAC | 3xFLAG-PatR fusion in pMBA131 |
| 58 | GTTTTATGCATAAAAACTCCTTGCGGCGGAGACAC | |
| 53 | GTTTTCTCGAGACTTCCGACCTGATGCCTTCGCCT | |
| 54 | GTTTTGAGCTCCTACTCCGTCGGCTTGGGTTTCTT | |

| | | |
|---|---|---|
| 978 | GTTTTATCGATCCAATAATGGGAACAACTGAAAC | *patR* promoter fused to *gfpmut2* |
| 979 | GTTTTCTCGAGCTCCTCCTAAAGTCATCACTTAC | |
| 49 | GTTTTCTCGAGATCTTCTTTTCTACCAAACACCAT | HetR-3xFLAG fusion plus downstream sequences in pMBA132 |
| 70 | GTTTTCTGCAGACCCTTATGACAAAGGACTTATAA | |
| 50 | GTTTTGAGCTCGCACCCAGAGTGAATAAAAGTACT | |
| 51 | GTTTTGAGCTCGCAGCCGTTGATTTTAACCAGTAT | |
| 47 | GTTTTGGATCCACCCTTATGACAAAGGACTTATAA | HetR-3xFLAG fusion plus downstream sequences in pCSRO |
| 52 | GTTTTGGATCCGCAGCCGTTGATTTTAACCAGTAT | |

Restriction sites are indicated underlined. T7 promoter is shown in bold.

**Table S5.** Plasmids.

| Name | Description |
|---|---|
| pCSRO | $Sm^RSp^R$, *sacB*-containing vector for conjugation of *Nostoc*[7] |
| pCSV3 | $Sm^RSp^R$, mobilizable vector for conjugation of *Nostoc*[8] |
| pRL278 | $Nm^R$, *sacB*-containing vector for conjugation of *Nostoc*[9] |
| pSAM301 | $Ap^RSm^RSp^R$, *nsiR1* promoter fused to *gfpmut2* and integration in alpha plasmid[5] |
| pSAM270 | $Ap^RSm^RSp^R$, *hetR* promoter fused to *gfpmut2* and integration in alpha plasmid[6] |
| pMBA20 | $Ap^RSm^RSp^R$, containing *rnpB* promoter and integration in alpha plasmid[10] |
| pMBA62 | $Sm^RSp^R$, pCSRO derivative containing a BamHI fragment corresponding to the *patR* region with *patR* gene deleted (this work). |
| pMBA66 | $Sm^RSp^R$, pMBA62 derivative containing $Nm^R$ gene inserted in the *patR* locus (this work). |
| pMBA79 | $Ap^RSm^RSp^R$, derivative of pMBA20 with *patR* expressed from the *rnpB* promoter (this work). |
| pMBA118 | $Ap^RSm^RSp^R$, *patR* promoter fused to *gfpmut2* for integration in alpha plasmid (this work). |
| pMBA131 | $Ap^RSm^RSp^R$, derivative of pMBA20 with 3xFLAG sequence fused to *sfgfp* expressed from the *rnpB* promoter (this work). |
| pMBA132 | $Ap^RSm^RSp^R$, derivative of pMBA20 with *sfgfp* fused to 3xFLAG sequence expressed from the *rnpB* promoter (this work). |
| pMBA140 | $Sm^RSp^R$, derivative of pMBA131 with 3xFLAG sequence fused to *sfgfp* expressed from the *patR* promoter (this work). |
| pMBA149 | $Sm^RSp^R$, derivative of pMBA140 with 3xFLAG sequence fused to *patR* expressed from the *patR* promoter (this work). |
| pMBA154 | $Sm^RSp^R$, derivative of pMBA132 with *hetR* fused to 3xFLAG sequence expressed from the *hetR* promoter (this work). |
| pMBA157 | $Sm^RSp^R$, derivative of pMBA154 with *hetR* fused to 3xFLAG sequence expressed from the *hetR* promoter and sequences downstream of *hetR* (this work). |
| pMBA160 | $Sm^RSp^R$, derivative of pCSRO with *hetR* fused to 3xFLAG sequence expressed from the *hetR* promoter and sequences downstream of *hetR* (this work). |
| pMBA166 | $Sm^RSp^R$, derivative of pMBA132 with *alr2809* fused to 3xFLAG sequence expressed from the *alr2809* promoter (this work). |
| pMBA167 | $Sm^RSp^R$, derivative of pCSV3 with *alr2809* fused to 3xFLAG sequence expressed from the *alr2809* promoter (this work). |
| pMBA170 | $Sm^RSp^R$, derivative of pMBA132 with *vipp1* fused to 3xFLAG sequence expressed from the *vipp1* promoter (this work). |
| pMBA171 | $Sm^RSp^R$, derivative of pCSV3 with *vipp1* fused to 3xFLAG sequence expressed from the *vipp1* promoter (this work). |
| pMBA177 | $Sm^RSp^R$, derivative of pMBA132 with *patU3* fused to 3xFLAG sequence (this work). |
| pMBA178 | $Sm^RSp^R$, derivative of pCSV3 with *patU3* fused to 3xFLAG sequence expressed after insertion in the chromosome from the native promoter (this work). |
| pMBA179 | $Sm^RSp^R$, derivative of pMBA132 with *prpA* fused to 3xFLAG sequence (this work). |
| pMBA180 | $Sm^RSp^R$, derivative of pCSV3 with *prpA* fused to 3xFLAG sequence expressed after insertion in the chromosome from the native promoter (this work). |
| pMBA182 | $Sm^RSp^R$, derivative of pMBA132 with *trpB* fused to 3xFLAG sequence (this work). |
| pMBA183 | $Sm^RSp^R$, derivative of pCSV3 with *trpB* fused to 3xFLAG sequence expressed after insertion in the chromosome from the native promoter (this work). |
| pMBA188 | $Sm^RSp^R$, derivative of pMBA132 with *rnpA* fused to 3xFLAG sequence (this work). |
| pMBA189 | $Sm^RSp^R$, derivative of pCSV3 with *rnpA* fused to 3xFLAG sequence expressed after insertion in the chromosome from the native promoter (this work). |
| pMBA211 | $Sm^RSp^R$, derivative of pMBA132 with *alr2890* fused to 3xFLAG sequence expressed from the *alr2890* promoter (this work). |
| pMBA213 | $Sm^RSp^R$, derivative of pCSV3 with *alr2890* fused to 3xFLAG sequence expressed from the *alr2890* promoter (this work). |

| | |
|---|---|
| pMB2 | $Sm^RSp^R$, derivative of pMBA132 with *asl3888* fused to 3xFLAG sequence expressed from the *asl3888* promoter (this work). |

#### Overview on Supplementary Data

**Supplementary Data 1.** Normalized distribution of the 2638 identified proteins in 18 analyzed fractions in each of the 6 gradients.

**Supplementary Data 2.** Mean of the normalized distribution of the 2638 proteins in 18 analyzed fractions for the control or RNase treated gradients.

**Supplementary Data 3.** Clustering analysis of the distribution of proteins in the control gradients.

**Supplementary Data 4.** Log2 foldchange and fdr values for the comparison between control and RNase treated gradients in each fraction for the 333 RNA-associated proteins.

**Supplementary Data 5.** Clustering analysis of the log2 foldchange data for the 333 shifting proteins.

**Supplementary Data 6.** Modified TriPepSVM prediction for the *Nostoc* proteome.

**Supplementary Data 7.** Elements of groups shown in the diagram of Figure 3d.

**Supplementary Data 8.** Modified TriPepSVM prediction for the *Nostoc* proteome and its homologs in 55 cyanobacterial proteomes.

**Supplementary Data 9.** ClustalW multiple alignment of the PatR homolog sequences in 107 selected cyanobacterial strains.
